# Localized Cardiolipin Synthesis is Required for the Assembly of MreB During the Polarized Cell Division of *Chlamydia trachomatis*

**DOI:** 10.1101/2022.01.12.476008

**Authors:** Scot P. Ouellette, Laura A. Fisher-Marvin, McKenna Harpring, Junghoon Lee, Elizabeth A. Rucks, John V. Cox

**Author notes:** = contributed equally (Laura A. Fisher-Marvin, McKenna Harpring, Junghoon Lee). Author order determined alphabetically. Corresponding authors: Scot P. Ouellette, Department of Pathology and Microbiology, and Biochemistrym, University of Nebraska Medical Center, Center 985900 Nebraska Medical Center, DRC2 5029, Omaha, NE 68198; John V. Cox, Department of Microbiology, Immunology, University of Tennessee Health Science, 858 Madison Ave, Memphis, TN 38163.

## Abstract

Pathogenic *Chlamydia* species are coccoid bacteria that use the rod-shape determining protein MreB to direct septal peptidoglycan synthesis during their polarized cell division process. How the site of polarized budding is determined in this bacterium, where contextual features like membrane curvature are seemingly identical, is unclear. We hypothesized that the accumulation of the phospholipid, cardiolipin (CL), in specific regions of the cell membrane induces localized membrane changes that trigger the recruitment of MreB to the site where the bud will arise. To test this, we ectopically expressed cardiolipin synthase (Cls) and observed a polar distribution for this enzyme in *Chlamydia trachomatis*. In early division intermediates, Cls was restricted to the bud site where MreB is localized and peptidoglycan synthesis is initiated. The localization profile of Cls throughout division mimicked the distribution of lipids that stain with NAO, a dye that labels CL. Treatment of *Chlamydia* with 3’,6-dinonylneamine (diNN), an antibiotic targeting CL-containing membrane domains, resulted in redistribution of Cls and NAO-staining phospholipids. In addition, MreB localization was altered by diNN treatment, suggesting an upstream regulatory role for CL-containing membranes in directing the assembly of MreB. This hypothesis is consistent with the observation that the clustered localization of Cls is not dependent upon MreB function or peptidoglycan synthesis. Furthermore, expression of a CL-binding protein at the inner membrane of *C. trachomatis* dramatically inhibited bacterial growth supporting the importance of CL in the division process. Our findings implicate a critical role for localized CL synthesis in driving MreB assembly at the bud site during the polarized cell division of *Chlamydia*.

## Introduction

The obligate intracellular bacterium *Chlamydia* undergoes a unique biphasic developmental cycle that alternates between a non-dividing infectious form, the elementary body or EB, and a non-infectious dividing form, the reticulate body or RB [1]. *Chlamydia* species such as *C. trachomatis* and *C. pneumoniae* are major pathogens of humans causing sexually transmitted infections and community acquired pneumonia, respectively [2–5]. Due to their obligate intracellular nature, an understanding of the underlying mechanisms these bacteria use to accomplish essential processes is lacking. However, the recent development of selected genetic tools for *Chlamydia* has facilitated studies on the basic microbiology of these unique bacteria [6–9].

*Chlamydia* has undergone significant genome reduction in evolving to obligate intracellular dependence [10]. Interestingly, even after genomic reduction, *Chlamydia,* a coccoid bacterium, encodes a number of rod-shape determining proteins including the actin-like protein, MreB [11–14]. One gene that has been lost in these bacteria is *ftsZ*, which encodes the tubulin-like FtsZ protein that orchestrates the cell division process in most bacteria [15]. Whereas most model bacteria divide by FtsZ-dependent binary fission, we have demonstrated that *Chlamydia* undergoes an MreB-dependent polarized cell division process [11,13,16]. This budding-like mechanism of cell division initiates with the synthesis of a patch of peptidoglycan at the pole of *Chlamydia trachomatis* where the budding daughter cell will arise. As the polarized division process proceeds and peptidoglycan deposition continues, the peptidoglycan structure can be visualized as a ring that is retained at the septum that forms between the budding daughter cell and the mother cell [13,17,18]. Over time, the bud is enlarged as the peptidoglycan ring grows until the budding daughter cell is equal in volume to the mother cell. At this point, constriction of the dividing cells occurs with concomitant loss of the peptidoglycan structure. Inhibitor studies have revealed that the activity of MreB, which overlaps the distribution of the septal peptidoglycan ring, is essential for the polarized cell division process of *Chlamydia* [11,13,18]. We have further demonstrated that chlamydial MreB has a unique N-terminal domain, as compared to other bacterial orthologs, that possesses membrane binding properties that may allow it to functionally substitute for the lack of FtsZ in these organisms [13]. However, the means by which MreB is recruited to the site of budding was unclear prior to the current study.

The Gram-negative pathogenic *Chlamydia* species are unusual in lacking a classical peptidoglycan sacculus to define their cell shape [18]. Their cell walls are essentially lipid-based and likely adopt the most thermodynamically favorable shape within an aqueous environment: that of a sphere. This morphology presents unique challenges for context-dependent protein localization as the inner leaflet of the inner membrane displays uniform negative membrane curvature. Given that MreB is the key driver of cell division in these bacteria and that cell division is initiated from one pole of the cell [11, 16], the uniform curvature of the inner membrane poses an additional conundrum as MreB is excluded from areas of negative membrane curvature in other bacteria [19–21]. In rod-shaped bacteria like *E. coli*, one characteristic of the cell poles is that they are enriched in anionic phospholipids (aPLs) like cardiolipin (CL) [22, 23]. CL is particularly suited for such sites as it is a conical phospholipid with a small head group and four acyl chains, and its clustering allows it to spontaneously introduce negative membrane curvature [24].

We hypothesized that localized CL synthesis in *Chlamydia* induces membrane bending that in turn enables MreB recruitment to initiate cell division. Alternatively, CL may directly recruit chlamydial MreB, perhaps through an interaction with its unique N-terminal domain [13].

Although CL is not a major component of the chlamydial cell membrane [25], a study from the Rock lab demonstrated that *Chlamydia* synthesizes CL and identified a candidate CL synthase (*cls*) gene, *ct284* [26]. To initiate our studies, we performed a bioinformatic analysis of chlamydial *cls* and monitored its transcription during the developmental cycle to determine that it is expressed during the stage when RBs are undergoing growth and division. Inducible expression of a 6xHis tagged Cls in *C. trachomatis* revealed that this enzyme accumulated at the bud site where MreB is localized and peptidoglycan synthesis occurs prior to the initiation of daughter cell outgrowth. As the budding process proceeded, Cls was maintained at the leading edge of the daughter cell, while MreB was retained at the septum between the daughter and the mother cell. The localization profile of Cls throughout the division process mimicked the distribution of lipids that stain with NAO, a dye that labels aPLs, including CL. Treatment of *Chlamydia* with 3’,6-dinonylneamine (diNN), an antibiotic that targets CL-containing membrane domains [27, 28], resulted in a redistribution of Cls and NAO staining aPLs. Importantly, MreB localization was significantly altered in diNN-treated cells suggesting an upstream regulatory role for CL-containing membranes in directing the assembly of MreB. This hypothesis is consistent with the observation that the clustered localization of Cls is not dependent upon MreB function or peptidoglycan synthesis. Furthermore, the expression of a mitochondrial derived CL-binding protein [29] at the inner membrane of *C. trachomatis* dramatically inhibited bacterial growth supporting the importance of CL in the division process. Our findings implicate polarized CL synthesis as an early event that is critical for the recruitment of MreB to the bud site during the polarized cell division process of *Chlamydia*.

## Results

### The cardiolipin synthase ortholog of C. trachomatis localizes to the leading edge of the dividing RB

In 2015, a chlamydial cardiolipin synthase was bioinformatically identified by Yao et al. but not further characterized [26]. This same study identified changes in CL content within infected cell cultures that were consistent with a chlamydial origin, concluding that *Chlamydia* synthesizes its own CL from the phosphatidic acid pool [26]. The gene designation for *cls* in *C. trachomatis* is *ct284/ctl0536*. The chlamydial *cls* is conserved in pathogenic *Chlamydia* species (Suppl. Fig. 1A), which lack a peptidoglycan sacculus, and weak homology (<50% similarity) was found to orthologous proteins in more distantly related *Chlamydia* genera like *Waddlia* and *Protochlamydia* that possess peptidoglycan saccula (data not shown). As an initial indicator of when Cls functions during the chlamydial developmental cycle [30], we monitored its transcription by RT-qPCR. Transcripts for *cls* were not reliably detected at the 0 and 1h post-infection (hpi) timepoints but increased from 3 to 16hpi before declining (Suppl. Fig. 1B). Peak transcription at 16hpi is consistent with the RB phase of the developmental cycle [12, 31], which suggests a potential function for Cls in RB growth and division.

To initiate our studies of the function of Ct284/Cls, we created a transformant of *C. trachomatis* L2 carrying a plasmid with an anhydrotetracycline (aTc)-inducible hexahistidine-tagged Cls (Cls_6xH). We then assessed the localization of the Cls_6xH protein in *Chlamydia*. After infecting cells with this transformant, we induced expression and fixed the cells at 10.5hpi, a time when they are undergoing their first division [16]. As we have described previously, the major outer membrane protein (MOMP) is highly polarized to one side of the cell at this stage of the developmental cycle, and the daughter cell arises from the MOMP-enriched pole of the mother cell [16]. For this and subsequent experiments, we have defined several intermediates in the chlamydial polarized division process. Pre-division intermediates (“round” cells) have no obvious outgrowth of the budding daughter cell. In “early” division intermediates the daughter cell is <15% of the mother cell volume, in “mid/late” division intermediates the daughter cell is between 15-80% of the mother cell volume, and in the “two-cell” stage of division the daughter cell is >80% of the mother cell volume. Cls_6xH accumulated at the MOMP-enriched side of the mother cell in round cells, and it remained at the leading edge of the budding daughter cell after division started in mid/late intermediates where it overlapped the distribution of MOMP (Fig. 1A). To determine whether this was a general feature of phospholipid synthases, we also monitored the localization of PsdD_6xH, which is an annotated phosphatidylethanolamine synthase. In contrast to Cls_6xH, PsdD_6xH localized on the opposite side from the MOMP-enriched pole of the cell both before and after division initiated (Fig. 1B). PsdD_6xH was only detected in the budding daughter cell at the two-cell stage of the first division (data not shown). Finally, we compared the localization pattern of these proteins to MreB_6xH, a characterized division protein in *Chlamydia* that substitutes for FtsZ [11, 14]. Prior to the initiation of division, MreB_6xH, like Cls_6xH, is clustered at a site on the MOMP-enriched side of the mother cell. However, at the onset of daughter cell growth, MreB_6xH is retained at the septum where it forms a ring structure beneath the growing bud (Fig. 1C) [13]. These data demonstrate that Cls and MreB both accumulate in polar clusters prior to the initiation of daughter cell growth. However, as the budding process initiates, Cls is maintained at the leading edge of the budding daughter cell, while MreB is retained at the septum between the daughter and the mother cell.

**Figure 1.**
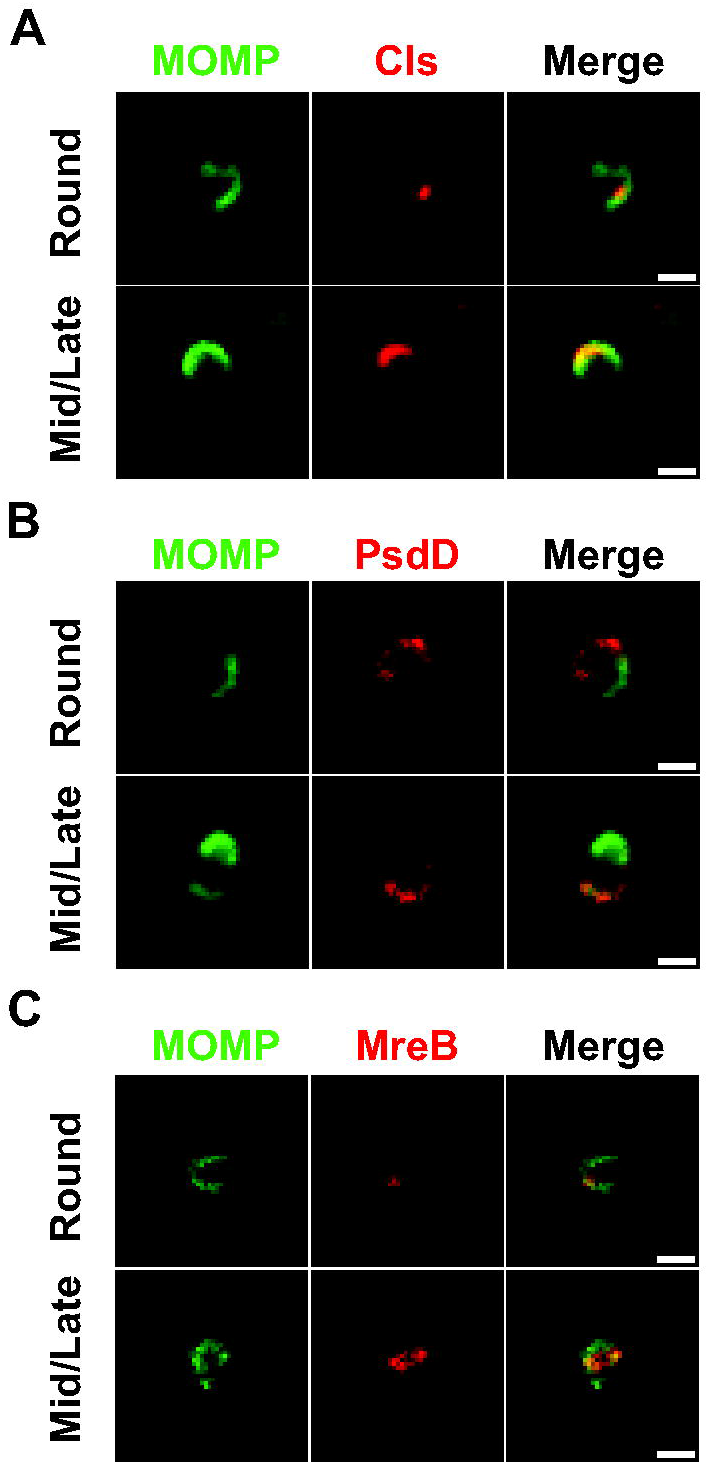
Localization of Cls and other markers during early and late stages of the initial RB division. McCoy cells were infected with *C. trachomatis* L2 transformants carrying plasmids encoding inducible constructs for (A) Cls_6xH, (B) PsdD_6xH, or (C) MreB_6xH. Expression of the construct was induced at 4hpi with 10nM anhydrotetracycline (aTc). At 10.5hpi, cells were fixed and processed for immunofluorescence analysis as described in Materials and Methods. Shown are representative images for each in both a pre-dividing cell (“Round”) and one in a “Mid/Late” division intermediate. Note the colocalization of Cls with the leading edge of the budding daughter cell whereas PsdD is at the opposite side in the mother cell. MreB forms a ring under the daughter cell within the division septum. Images were acquired on a Nikon Ti2 spinning disc confocal microscope using a 60X lens objective. Images are representative of at least three independent experiments. Scalebar = 2 μm.

To investigate the potential function of Cls at later stages of the developmental cycle, cells were infected with the Cls_6xH transformant, the expression of the fusion was induced at 8h post-infection (hpi) with different concentrations of aTc, and the cells were fixed and processed for immunofluorescence analysis (IFA) at 24hpi. The expression of Cls_6xH had no obvious effect on inclusion size and the protein exhibited a polar pattern of localization in individual cells within the inclusion (Fig. 2A). Overexpression of the wild-type Cls isoform did display a statistically significant but modest negative impact on growth of the organism as assessed by inclusion forming unit assay (a chlamydial equivalent to colony forming unit assay). After induction of Cls_6xH with 5nM aTc, chlamydial IFUs were 23% +/-3% (p<0.001) of the uninduced, untreated control levels.

**Figure 2.**
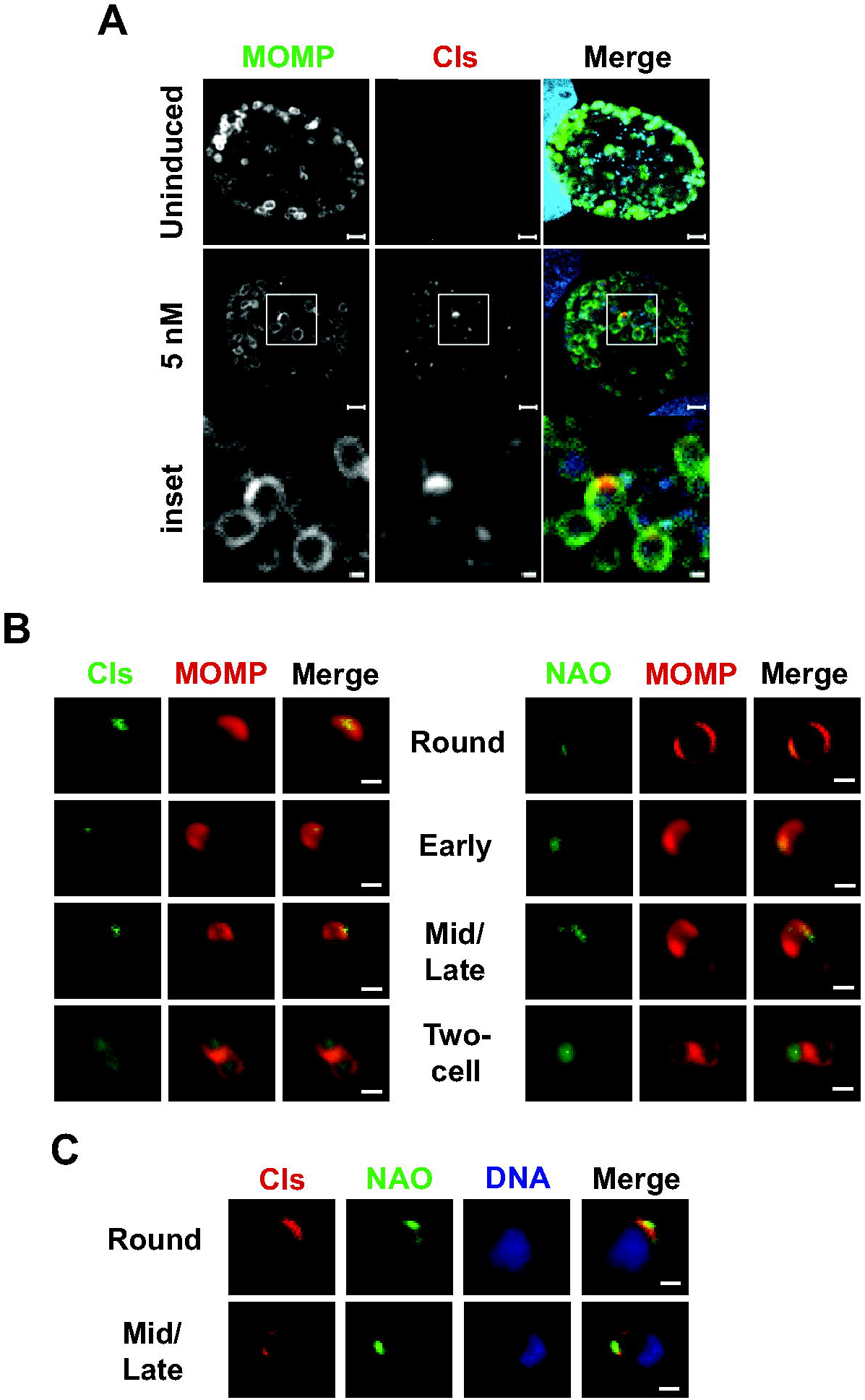
Localization of Cls_6xH at later stages of the developmental cycle and co-localization with anionic phospholipids (aPLs) as assessed by NAO staining. (A) Localization of wild-type Cls_6xH in *C. trachomatis* L2. Cells were infected, induced or not for expression at 4hpi with 5nM anhydrotetracycline (aTc), and samples were collected and processed at 24hpi. Bacteria were labeled with an antibody targeting MOMP (major outer membrane protein - green), and the construct was labeled with an antibody against the 6xH tag (red). Nuclear DNA was labeled with DAPI and is visualized within the merged image only in the blue channel. The boxed region within each induced image is shown below as an enlarged inset. Note each isoform shows a discrete localization pattern to one side of the bacteria. Images were acquired on a Zeiss AxioImager.Z2 equipped with an Apotome2 using a 100X lens objective. Images are representative of at least three independent experiments. Scalebar of full inclusion images = 2 μm Scalebar of inset = 0.5 μm (B) Localization of Cls and aPLs in partially purified RBs. HeLa cells were infected with the Cls_6xH transformant, and expression of the construct was induced at 20hpi with 10nM aTc. RBs were isolated and partially purified from infected cells at 22hpi as described in Materials and Methods prior to imaging the distribution of MOMP, Cls, or aPLs (using 250nM NAO). The localization of each marker is shown at Round, Early, Mid/Late, and Two-cell stages of the budding division process. (C) Co-localization of Cls with NAO-stained aPLs. RBs were induced for expression of Cls_6xH and partially purified as described above. RBs were then imaged for Cls, aPLs (using NAO), and DNA (using Hoechst 33342) at Round and Mid/Late stages of division. Images for (B) and (C) were acquired with a Zeiss AxioImager2 microscope equipped with a 100x oil immersion PlanApochromat lens. Scalebar = 2 μm

Since it was difficult to assess the stage of division of cells within the inclusion at the 24h time point, we induced the expression of Cls_6xH with aTc at 20hpi and partially purified *Chlamydia* from infected HeLa cells at 22hpi when the majority of bacteria within the inclusion are dividing RBs. IFA of these cells revealed that Cls_6xH localization in dividing RBs at this stage of the developmental cycle was very similar to that observed in cells undergoing their first division (Fig. 1A). Cls_6xH accumulated in a polar cluster in round cells and at the leading edge of budding daughter cells in early and mid/late budding intermediates (Fig. 2B). At the two-cell stage, the localization of Cls_6xH became more diffuse within the daughter and mother cells (Fig. 2B). Identical results were obtained when infected cells were fixed prior to partially purifying *Chlamydia*, indicating that the profiles observed were not a consequence of the isolation procedure (data not shown). This approach also enabled us to visualize the distribution of anionic phospholipids (aPLs) in bacteria at various stages of polarized division. Nonyl acridine orange (NAO) binds the aPLs CL and phosphatidylglycerol, and this dye has been used to visualize the distribution of these lipids in other bacteria [22, 23]. Attempts to use NAO to determine whether the polar Cls_6xH staining observed in *Chlamydia* at 24hpi (Fig. 2A) corresponded to regions of the chlamydial membrane enriched in these aPLs were unsuccessful because of the high levels of background staining observed in infected cells stained with this dye (data not shown). However, when we stained partially purified *Chlamydia*, in which the expression of Cls_6xH was induced, with NAO, the distribution of aPLs, like Cls_6xH, was primarily in a polar cluster in round cells. NAO labeling was restricted to the leading edge of budding daughter cells in early and mid/late division intermediates (Fig. 2B). At the two-cell stage, Cls_6xH became more diffuse (Fig. 2B). There was no detectable NAO staining observed in uninduced cells under the conditions used in our assays (data not shown), and the images in Figure 2C illustrate that Cls_6xH overlapped the distribution of NAO-staining phospholipids in induced cells undergoing division. These data indicate that the expression of Cls_6xH induces the accumulation of aPLs at the leading edge of budding daughter cells during polarized division.

### The transmembrane domain of Ct284/Cls is necessary for its restricted localization

Interestingly, our bioinformatics analyses revealed that the transmembrane (TM) domain region at the N-terminus of the chlamydial Cls is notably shorter than the TM domains of *E. coli* or *B. subtilis* Cls orthologs. Bacterial cardiolipin synthases typically possess two TM domains such that the enzymatic domain is oriented towards the cytosol [32, 33]. TM prediction programs confirmed that the chlamydial Cls possesses only a single predicted TM domain (data not shown).

To determine whether the TM domain of Cls was sufficient to direct its restricted membrane localization, we created a chlamydial transformation plasmid encoding an aTc-inducible construct where the chlamydial Cls TM domain was fused to GFP (Cls_TM_GFP). This plasmid also constitutively expresses mCherry. Transformants of *C. trachomatis* serovar L2 were obtained, and the fusion protein was expressed after infecting a monolayer of cells. As seen in Figure 3A, the GFP signal was detected and localized to discrete sites on the RB membrane in live-cell images. These data indicate that the TM domain of Cls is sufficient to direct its clustered localization.

**Figure 3.**
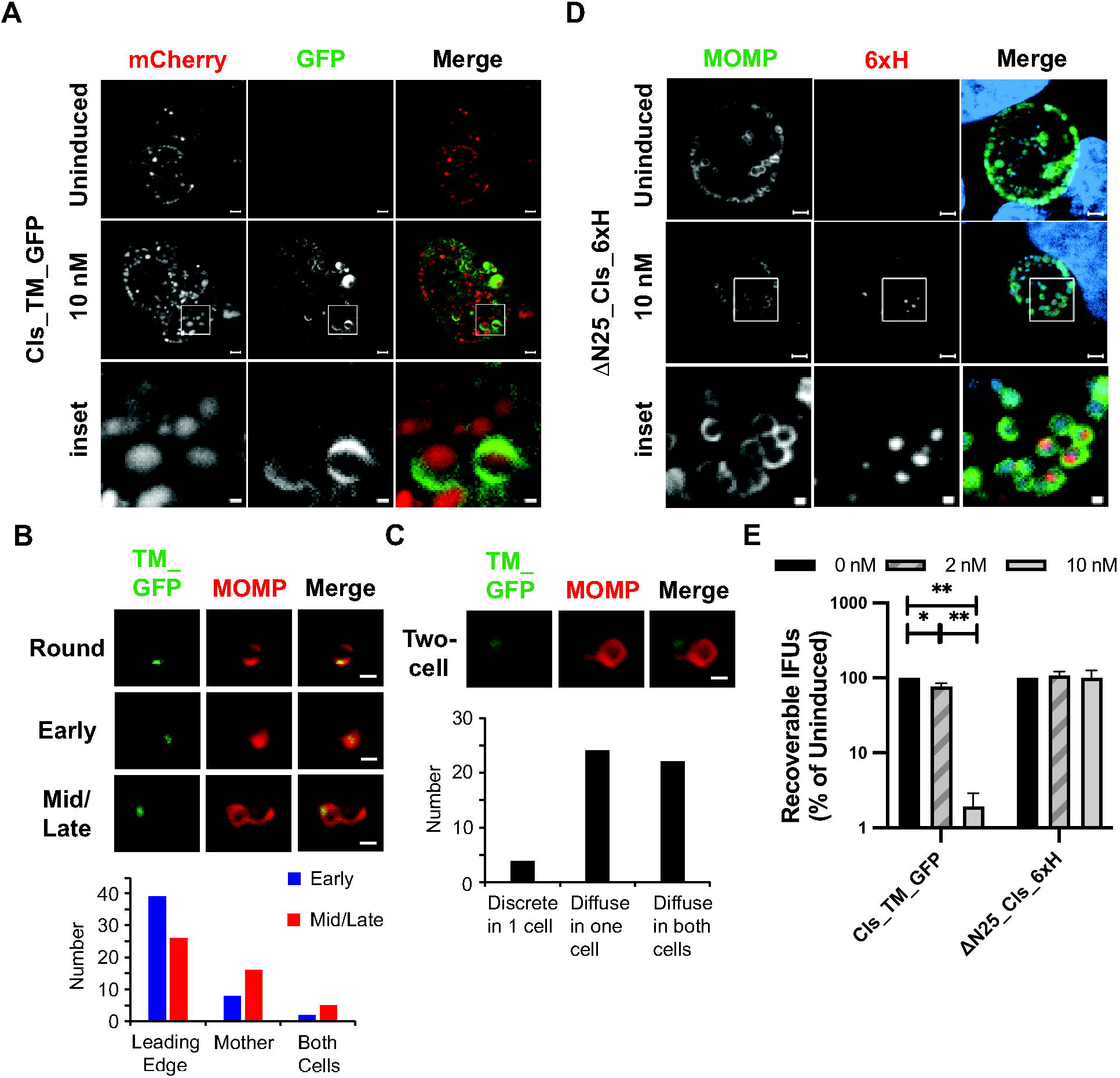
Functional evaluation of the Cls transmembrane (TM) domain on protein localization. (A) *C. trachomatis* L2 was transformed with an anhydrotetracycline (aTc) inducible vector encoding the Cls TM domain (first 25 residues) fused to GFP; mCherry is encoded on the plasmid backbone to be constitutively expressed. Cells were infected with this strain, and expression of Cls_TM_GFP was induced or not at 16hpi with 10nM aTc. Localization of the marker and mCherry were acquired from live-cell imaging. (B) The localization of Cls_TM_GFP at Round, Early, and Mid/Late stages of division or (C) the two-cell stage of division was imaged and quantified from partially purified RBs. For (B) Cls_TM_GFP was localized to a polar cluster in round cells and primarily at the leading edge of the budding daughter cells, whereas for (C) it was localized as a discrete patch in one cell or diffuse in one or both cells. At least 50 cells were counted for each stage of division. Images were acquired with a Zeiss AxioImager2 microscope equipped with a 100x oil immersion PlanApochromat lens. Scalebar in panels B and C = 2 μm (D) Cells were infected with *C. trachomatis* L2 transformed with an aTc-inducible vector encoding Cls_6xH lacking the TM domain (i.e. starting at residue 26). Expression of the construct was induced or not at 4hpi, and samples were fixed and processed at 24hpi as described in the legend to Figure 2. For (A) and (D) the boxed region within each induced image is shown below as an enlarged inset. Note the discrete localization pattern to one side of the bacterial membrane in (A) but not in (D). Images were acquired on a Zeiss AxioImager.Z2 equipped with an Apotome2 using a 100X lens objective. Images are representative of at least three independent experiments. Scalebar of full inclusion images = 2 μm Scalebar of inset = 0.5 μm (E) Cells were infected with the indicated transformants and processed as described in the legend to Figure 3 to quantify IFU production during the primary infection. For each transformant, the uninduced values were arbitrarily set to 100%, and the effect of overexpression at either 2nM or 10nM aTc when added at 4hpi is expressed as a percentage of the wild-type. Data are the average of three independent experiments performed in triplicate. ** = p<0.001

Since it was again difficult to assess the distribution of Cls_TM_GFP during specific stages of division when infected cells were analyzed (Fig. 3A), we induced the expression of this fusion by adding aTc to partially purified *Chlamydia* transformant grown in axenic media (Optimem supplemented with 1mM glucose-6-phosphate and 1mM glutamine [34]). IFA analyses of these cells and quantification of the localization pattern revealed that Cls_TM_GFP exhibited a localization profile indistinguishable from wild-type Cls, as it accumulated in a polar cluster in rounds cells and at the leading edge of budding daughter cells at early and mid/late stages of division (Fig. 3B). At the two-cell stage, the localization of Cls_TM_GFP became more diffuse within the mother cell or both cells (Fig. 3C).

Given the discrete localization of GFP when fused to the Cls TM domain and the discrete localization of full-length Cls_6xH (Figs. 1&2), we next determined if the TM domain was critical for the clustered localization of Cls. To test this, we created an N-terminal truncation of Cls_6xH lacking its TM domain (Δ25Cls_6xH) and expressed this in *Chlamydia*. As seen in Figure 3D, Δ25Cls_6xH localized to the cytosol of the bacteria. We conclude from these data that the TM domain of the chlamydial Cls directs its restricted localization to specific regions of the inner membrane.

Finally, we determined the impact of overexpression of the Cls_TM_GFP and ⊗25Cls_6xH constructs on chlamydial growth using the IFU assay (Fig. 3E). Infected cells were treated with aTc or not at 8hpi using two different concentrations of inducer. We did not measure any growth defects from overexpressing the Δ25Cls_6xH construct but did note that extended overexpression of Cls_TM_GFP with 10nM, but not 2nM, aTc resulted in abnormally enlarged bacteria (Suppl. Fig. 2) and fewer infectious progeny (Fig. 3E), suggesting that accumulation of the Cls_TM domain in the inner membrane may alter membrane organization and disrupt normal cell division.

### Cls_6xH localization is dependent upon its interaction with CL-containing membrane microdomains

3’,6-dinonylneamine (diNN) is an amphiphilic aminoglycoside that interacts with aPLs [27, 28]. Treatment of *P. aeruginosa* with this drug induced the redistribution of NAO-staining phospholipids from the poles to the sidewall of the cell and affected the permeability and morphology of this organism [35]. Concentrations of diNN that were lower than that required to alter lipid localization and cell growth of *P. aeruginosa* were toxic to HeLa cells (data not shown), so the effect of the drug on Cls_6xH localization could not be monitored by treating *Chlamydia*-infected cells. To circumvent this, we monitored the effect of diNN on partially purified *Chlamydia* transformants isolated from infected cells at 22hpi.

When the expression of Cls_6xH was induced in partially purified cells grown in axenic media containing aTc for 1.5hrs, Cls_6xH and NAO-staining phospholipids were primarily restricted to the leading edge of budding daughter cells in polarized division intermediates (Fig. 4A), similar to the result obtained when the expression of the fusion was induced during infection (Fig. 2B). Quantification of the staining profiles observed in cells grown in axenic media (Fig. 4B) revealed that both Cls_6xH and NAO were in a polar cluster in rounds cells and present at the leading edge of the daughter cell in most cells during early and mid/late stages of division. The other observed localization profiles primarily reflected the distribution of Cls_6xH and NAO staining during the two-cell stage prior to cell constriction and separation. To assess the effect of diNN on Cls_6xH and aPL localization, cells were incubated in axenic media containing aTc and 5μM diNN for 1.5hrs. In most of the cells, this treatment resulted in a dramatic and statistically significant redistribution of both Cls_6xH and NAO staining to a discrete region in the mother cell membrane (Fig. 4A&B). In a subset of the early division intermediates treated with diNN, the distribution of Cls_6xH and NAO-staining phospholipids was similar to that seen in untreated controls. Importantly, there was no statistically significant difference between the Cls and NAO localization profiles at the different stages and between treatments. The molecular basis for the differential effect of diNN on the localization of Cls_6xH and NAO staining phospholipids in cells at varying stages of division is unclear at this time.

**Figure 4.**
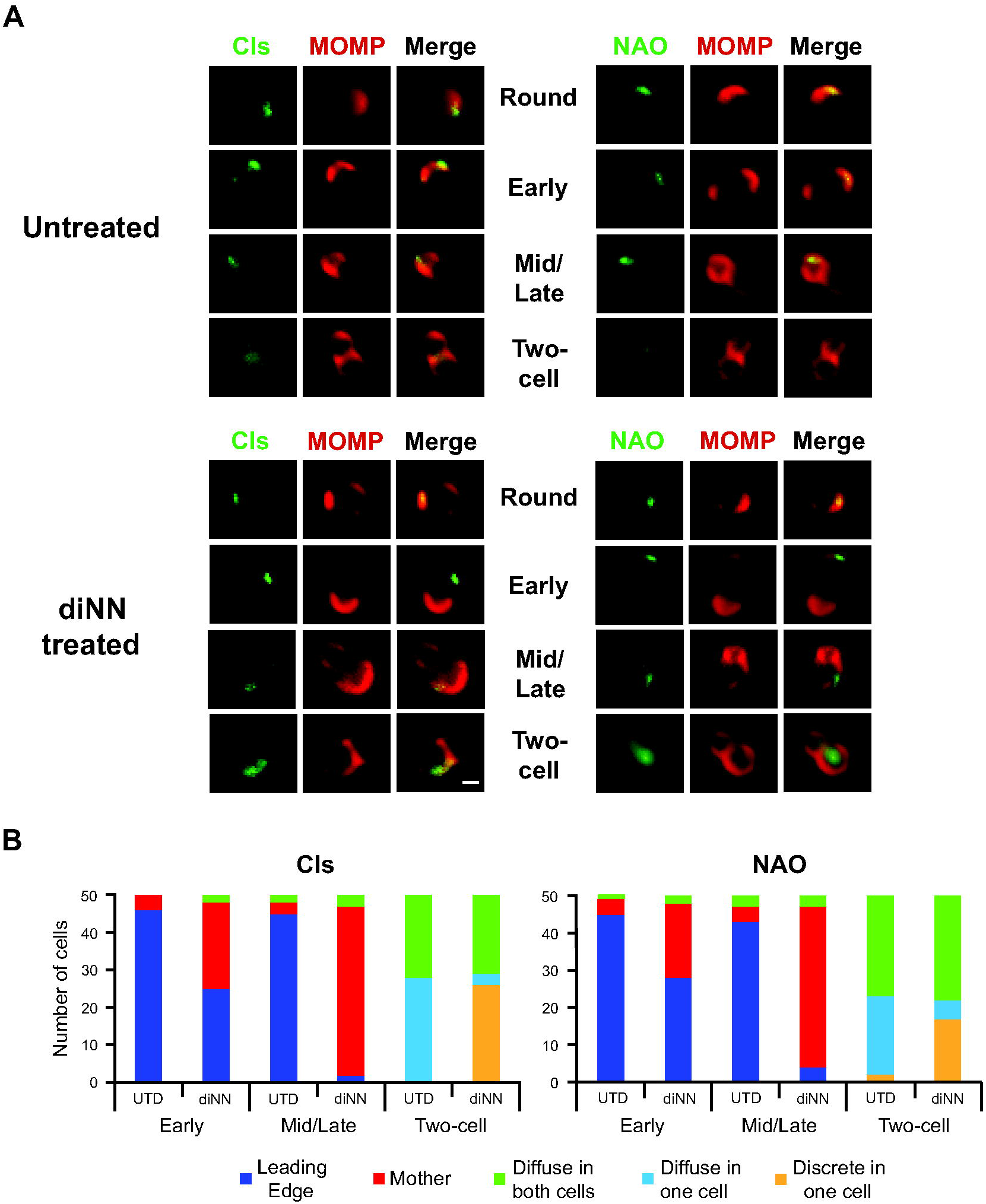
Effect of the CL-targeting antibiotic 3’,6-dinonylneamine (diNN) on localization of Cls and aPLs at different stages of the division process. HeLa cells were infected with the Cls_6xH transformant, and, at 22hpi, RBs were isolated and partially purified. Cls_6xH expression was induced by incubating cells in axenic media with 10nM aTc in the presence or absence of 5 μM diNN, and cells were fixed after 1.5h. Samples were processed for the major outer membrane protein (MOMP), and Cls or NAO (using 250nM). (A) Representative images of the indicated markers in Untreated or diNN treated bacteria at the indicated stages of cell division. Images were acquired with a Zeiss AxioImager2 microscope equipped with a 100x oil immersion PlanApochromat lens and are representative of two independent experiments. Scalebar = 2 μm (B) 50 individual cells from the indicated stages of division from untreated and diNN-treated cultures were assessed for their Cls and NAO localization profiles. Localization profiles were categorized into leading edge of the budding daughter cell, diffuse in mother cell, diffuse in one cell, diffuse in both cells, or discrete in one cell.The differences in localization of Cls and NAO between treatment conditions at each stage of division were statistically analyzed using a chi-squared test to reveal that the changes observed during diNN treatment were statistically significant (p<0.001) for each division intermediate compared to untreated (UTD). There was no statistical difference between Cls and NAO localization for any condition tested.

In addition, the response of Cls_TM_GFP to diNN treatment was virtually identical to Cls_6xH, as Cls_TM_GFP redistributed to a discrete region in the mother cell membrane when the drug was included in axenic media during induction (Suppl. Figs. 3A&B). Like Cls_6xH, Cls_TM_GFP was retained in the leading edge of budding daughter cells in a subset of the early division intermediates treated with the drug (Suppl. Figs. 3A&B). Taken together these results indicate that the TM domain is sufficient to direct Cls to the leading edge of budding cells, and its restricted localization is dependent upon its association with CL-rich membrane microdomains.

To determine whether diNN has a general effect on the localization of phospholipid synthases in *Chlamydia*, we characterized the distribution of PsdD_6xH induced in axenic culture in the absence and presence of diNN. Similar to the results obtained at 10.5hpi (Fig. 1B), PsdD_6xH primarily accumulated in the mother cells in polarized division intermediates, and diNN had no apparent effect on its localization profile (Suppl. Fig. 4). These results strongly suggest that the accumulation of Cls_6xH at the leading edge of budding cells is at least in part dependent upon its interaction with aPLs via its transmembrane domain, and they illustrate a clear difference in the role of aPLs in regulating the localization of Cls and PsdD.

### Ectopic expression of a CL/phosphatidic acid binding protein in Chlamydia severely disrupts organism growth and morphology

As we could not treat infected cells with diNN to determine effects of disrupting CL on chlamydial growth and morphology, we looked to circumvent this limitation by overexpressing a CL-binding protein in *Chlamydia* with the hypothesis that this would interfere with normal CL function. In 2015, a small mitochondrial protein, C11orf83, was characterized as a CL-binding protein with an N-terminal membrane anchor [29]. Although C11orf83 also has affinity for phosphatidic acid (PA) and sulfatide, there is no evidence that *Chlamydia* has sulfatide in its membranes [25, 26], and PA is typically short-lived (in *E. coli*), and rapidly incorporated into other phospholipids [36, 37]. We constructed several codon-optimized C11orf83 variants for aTc-inducible expression in *Chlamydia*: C11orf83_6xH, Cls_TM_c11orf83_6xH (appending the Cls TM domain to the full length c11orf83), and TM_c11orf83_6xH (replacing the mitochondrial TM domain with a TM domain from *E. coli* OppB to target C11orf83_6xH to the periplasm). We previously validated the use of the *E. coli* OppB first TM domain to target a protein into the periplasm [38]. The predicted localization of these constructs is depicted in Figure 5A.

**Figure 5.**
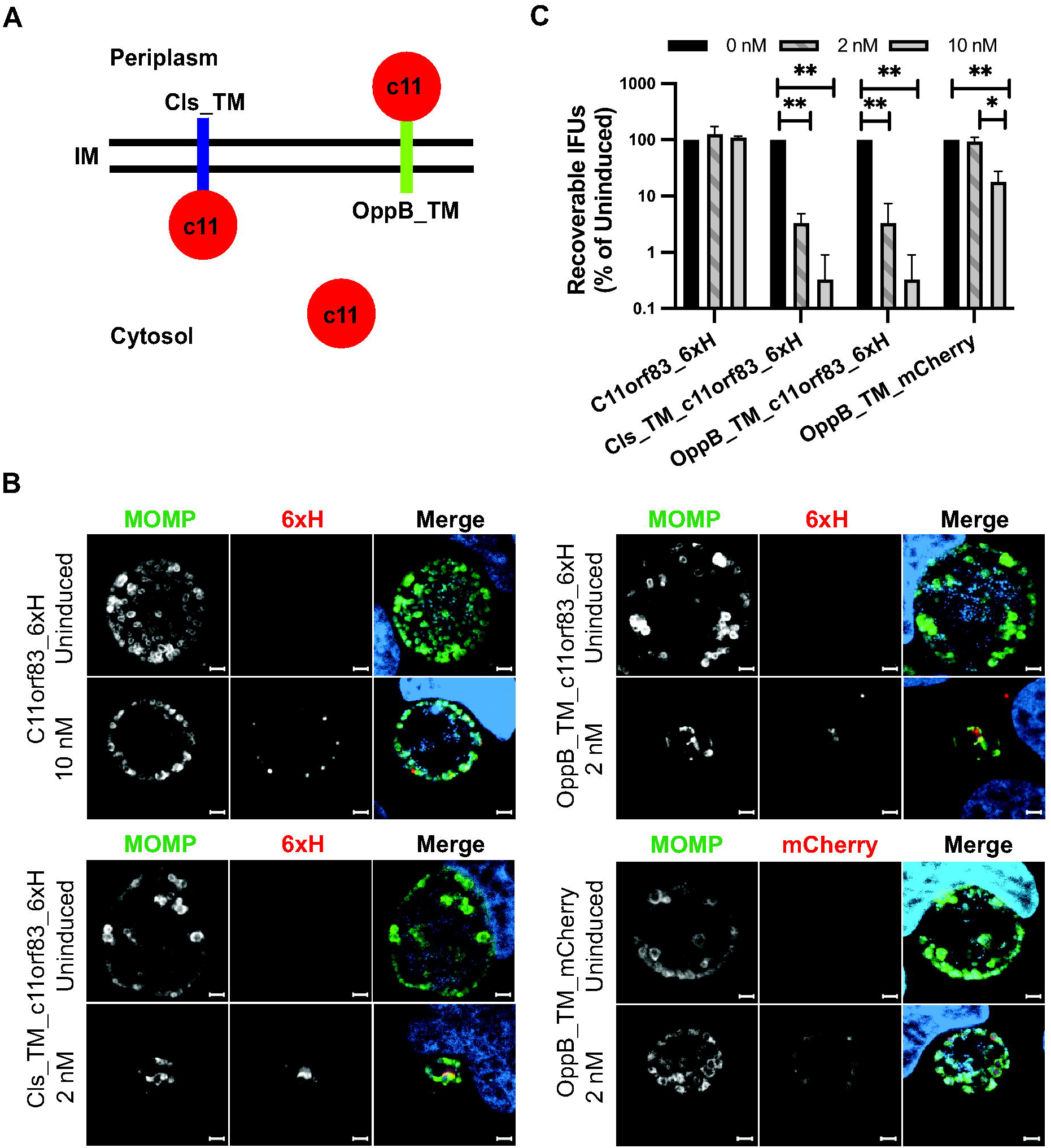
Effect on chlamydial growth and morphology of overexpression of a cardiolipin binding protein, C11orf83. (A) Schematic of the expected localization of each C11orf83 (c11) construct when expressed in chlamydiae. IM = inner membrane Cls_TM = first 25 residues of Cls fused to c11 OppB_TM = first TM domain of *E. coli* OppB fused to c11 (or mCherry). (B) Cells were infected with *C. trachomatis* L2 transformants carrying anhydrotetracycline (aTc) inducible plasmids and encoding the indicated constructs. Expression of each was induced or not at 8hpi with 2 or 10nM aTc as indicated, and cells were fixed and processed for immunofluorescence at 24hpi as described in the legend to Figure 2. Images were acquired on a Zeiss AxioImager.Z2 equipped with an Apotome2 using a 100X lens objective. Images are representative of at least three independent experiments. Scalebar = 2 μm (C) Cells were infected with the indicated transformants and processed as described in the legend to Figure 2 to quantify IFU production during the primary infection. For each transformant, the uninduced values were arbitrarily set to 100%, and the effect of overexpression at either 2nM or 10nM aTc when added at 4hpi is expressed as a percentage of the wild-type. Data are the average of three independent experiments performed in triplicate. * = p<0.05 ** = p<0.001

We infected cells with these transformants and induced the expression of the fusions at 8hpi with aTc, and then fixed and processed the cells for IFA at 24hpi. Although full-length C11orf83_6xH appeared polar in a subset of the cells, it accumulated in the cytosol (Fig. 5B), indicating that the mitochondrial TM domain was not capable of inserting into the chlamydial inner membrane. No obvious effects were noted on inclusion size or bacterial morphology when overexpressing this construct with 10nM aTc. In contrast, when this protein was tethered to the cytosolic facing side of the inner membrane with the Cls_TM domain, a dramatic reduction in chlamydial inclusion size was observed even when induced with a low concentration of aTc (2nM), and this was correlated with reduced bacterial numbers and abnormal bacterial cell shape. Note that this effect was not observed when the Cls_TM_GFP expression was induced with 2nM aTc (Fig. 3), indicating a specific effect of the C11orf83 at the membrane. Similar effects on inclusion size, bacterial numbers, and cell shape were observed when C11orf83_6xH was expressed in the periplasm and tethered to the outer leaflet of the inner membrane using the first TM domain from OppB of *E. coli* (Fig. 5B). As a control for overexpressing a protein in the periplasm, we expressed mCherry in the periplasm using the same OppB TM domain and observed its effects. In contrast to OppB_TM_c11orf83_6xH, the expression of OppB_TM_mCherry in the periplasm did not cause a noticeable decrease in inclusion size, and it did not appear to impact bacterial numbers.

To quantify the growth effects related to overexpression of the C11orf83 constructs or periplasmic mCherry, we performed an IFU assay to measure EB production from the various induced and uninduced cultures. As seen in Figure 5C, and consistent with our immunofluorescence analysis, overexpression of C11orf83_6xH in the cytosol or mCherry in the periplasm had minimal or no effect on chlamydial growth. In contrast, when C11orf83_6xH was tethered to either face of the inner membrane, a significant and dramatic decrease in IFUs was measured. Collectively, these data indicate that expression of the CL/PA-binding protein in the inner or outer leaflet of the chlamydial inner membrane is sufficient to disrupt chlamydial cell division.

### Cls_6xH localization is not disrupted by MreB or peptidoglycan synthesis inhibitors

Given the hypothesized role of Cls in cell division, we attempted to determine interactions between it and known cell division proteins using a bacterial adenylate cyclase-based two hybrid system (BACTH) [38, 39]. However, we were unable to detect interactions between Cls and any chlamydial cell division (e.g. Fts proteins [31]) or peptidoglycan synthesis-associated proteins (e.g. PBP, Mur proteins [12]; data not shown) we tested. Therefore, to determine an epistatic relationship between Cls, MreB, and peptidoglycan synthesis, more generally, we performed a series of experiments leveraging the known effects of A22, an MreB inhibitor [40], and D-cycloserine (DCS), which blocks early steps in the synthesis of peptidoglycan precursors [41], on the localization of these markers. We previously used a similar strategy to demonstrate that MreB acts epistatically to penicillin binding proteins in *Chlamydia* [11].

To perform these experiments, cells were infected with chlamydial transformants carrying inducible expression constructs of Cls_6xH or MreB_6xH. Expression of the constructs was induced at 8hpi with aTc. DCS was added at 2hpi whereas A22 was added at 16hpi. Cells were labeled for peptidoglycan using the Click-iT modifiable EDA-DA prior to fixation at 18hpi and processing for immunofluorescence [42]. In induced samples without antibiotic treatment, the distinct polar localization of Cls_6xH was observed, and peptidoglycan rings and other peptidoglycan intermediates were detected (Fig. 6A). However, these structures were generally not colocalized. MreB_6xH localization was consistent with prior studies, and the peptidoglycan labeling colocalized with MreB (Fig. 6B). Interestingly, with A22 treatment, MreB_6xH localization was significantly more diffuse and peptidoglycan labeling was lost, consistent with earlier findings [13], but Cls_6xH remained localized at a discrete site on the membrane. DCS, while grossly altering bacterial cell shape and the distribution of peptidoglycan, did not affect the clustered membrane localization of Cls_6xH or MreB_6xH. These data indicate that clustered localization of Cls is not impacted by the inhibition of MreB or peptidoglycan synthesis.

**Figure 6.**
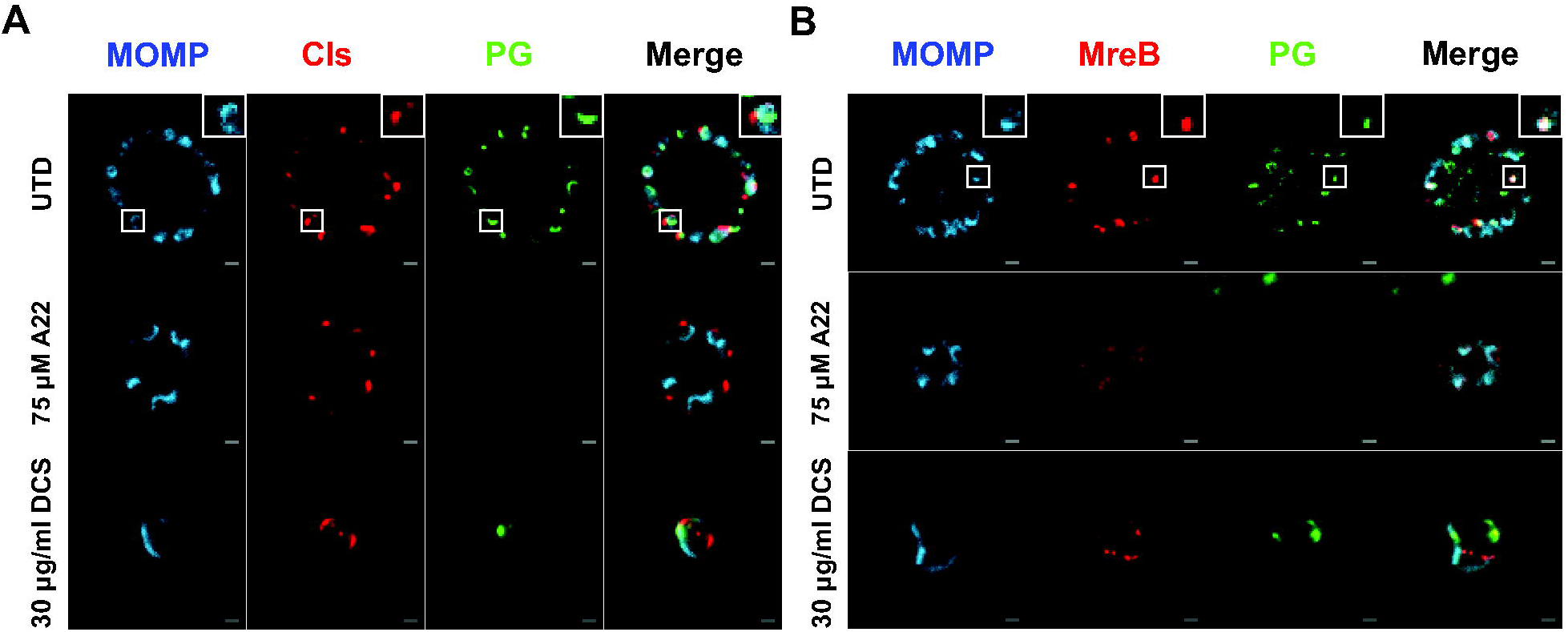
Effect of cell division inhibitors on localization of (A) Cls_6xH and (B) MreB_6xH. Cells were infected with transformants carrying anhydrotetracycline (aTc) inducible vectors encoding the indicated constructs. Expression of Cls_6xH or MreB_6xH was induced at 16 hpi, and cells were fixed and processed for immunofluorescence at 20 hpi. To visualize peptidoglycan (PG), cells were incubated with EDA-DA during the infection and processed using click-iT reagents as described in the Materials and Methods. To assess effects of disrupting the central coordinator of chlamydial cell division, MreB, the antibiotic A22 was added at 18hpi whereas peptidoglycan synthesis was disrupted using D-cycloserine (D-CS) added at 4hpi. Note the continued presence of Cls_6xH at a discrete site on the bacteria under all conditions tested. The insets within the upper row images are a zoomed-in view of the boxed regions with the larger image. Images were acquired on a Zeiss AxioImager.Z2 equipped with an Apotome2 using a 100X lens objective. Images are representative of at least three independent experiments. Scalebar = 2 μm

One interpretation of our data could be that premade Cls_6xH is recruited to the membrane by MreB, but its continued presence there is not dependent on MreB. We tested this possibility by pre-treating cells with A22 to disrupt MreB localization prior to inducing expression of Cls_6xH. However, under these conditions, induced Cls_6xH was still recruited to a discrete location on the membrane (Suppl. Fig. 5), indicating that its clustered distribution in cells occurs independently of MreB.

### Assembly of MreB is Altered in diNN-treated Chlamydia trachomatis

To directly investigate our hypothesis that membrane sub-domains enriched in CL are involved in directing MreB recruitment to specific sites in dividing *Chlamydia*, we investigated the effect of diNN on MreB localization in partially purified cells incubated in axenic media. A *Chlamydia* transformant carrying an aTc-inducible MreB_6xH plasmid was partially purified from infected cells at 22hpi, and the expression of MreB_6xH was induced by adding aTc to these cells grown in axenic media for 1 hr. The effect of diNN on MreB assembly was determined by pretreating purified cells with 5μM diNN for 30 minutes then inducing MreB expression for 1hr in the continued presence of the drug. In addition, to eliminate any effects related to the preexisting endogenous pool of MreB recruiting exogenously expressed MreB_6xH, purified cells were pretreated cells with 5μM diNN+ 75μM A22 for 30 minutes prior to inducing MreB_6xH expression for 1 hour in the presence of diNN only. For all treatment conditions, we quantified the number of dividing versus non-dividing cells in the population and observed that A22 pretreatment significantly reduced the overall number of dividing cells (Fig. 7A), consistent with the central role of MreB in orchestrating the synthesis of PG in the septum to drive daughter cell outgrowth [13]. Conversely, diNN treatment did not significantly impact the ratio of non-dividing to dividing cells (Fig. 7A).

**Figure 7.**
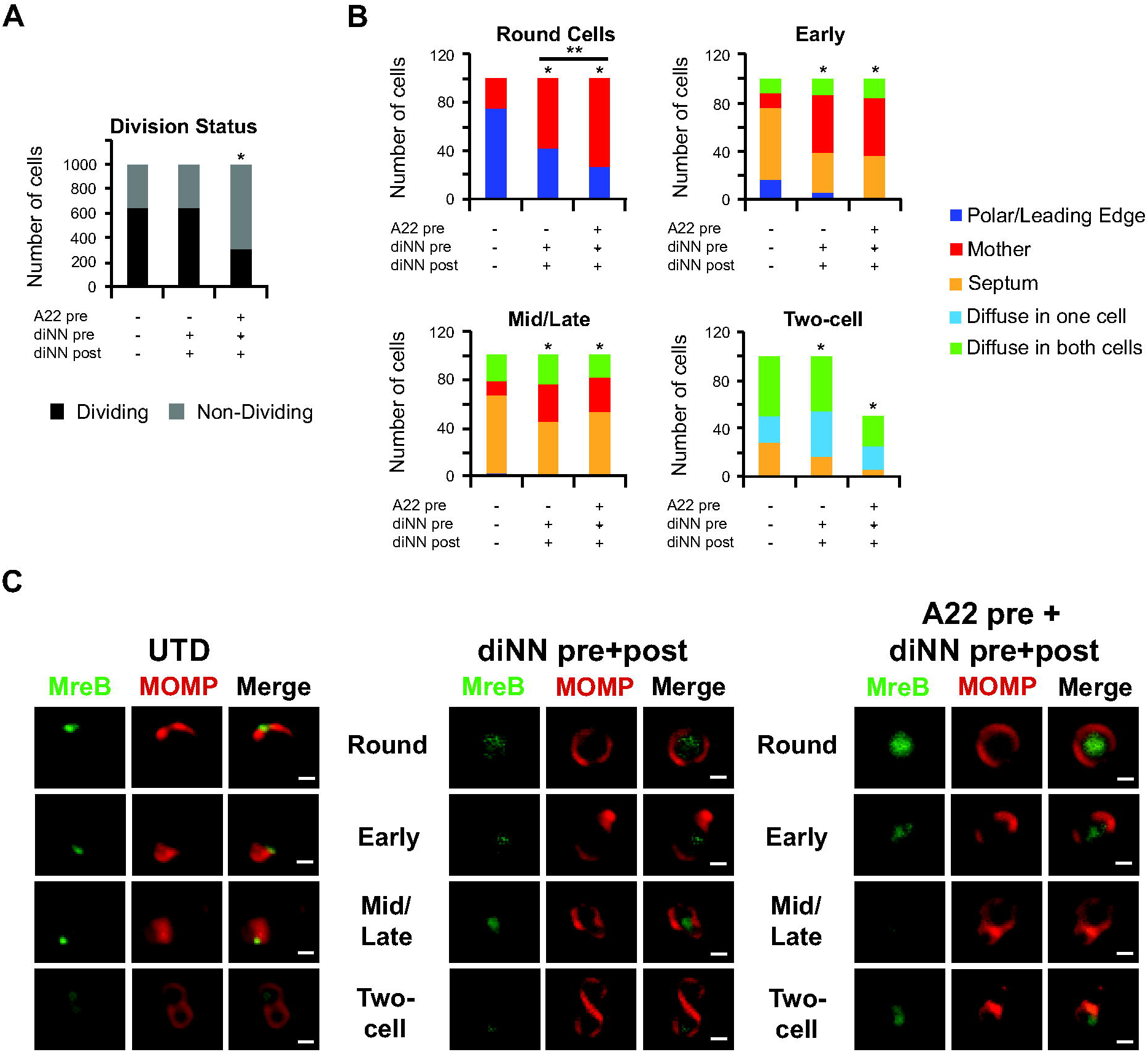
Effect of the CL-targeting antibiotic 3’,6-dinonylneamine (diNN) and the MreB-targeting antibiotic A22 on localization of MreB_6xH. HeLa cells were infected with the MreB_6xH transformant, and, at 22hpi, RBs were partially purified and MreB_6xH was induced by incubating the cells in axenic media containing 10nM aTc. In some instances, cells were preincubated with 5μM diNN or 5μM diNN + 75 μM A22 for 30 minutes and then induced with aTc in the presence of 5μM diNN alone. Following induction, the localization of MreB_6xH at various stages of division was assessed by staining cells with MOMP and 6xHis antibodies. (A) The total number of dividing versus non-dividing cells from the indicated culture conditions was quantified from 1000 total bacteria. * = p<0.0001 compared to the untreated (UTD) control as measured by chi-squared test. (B) The localization of MreB_6xH was assessed in individual cells from each stage of division from untreated cultures or cultures treated as indicated in the figure. Localization profiles were categorized into leading edge of the budding daughter cell/polar, diffuse in mother cell, diffuse in one cell, diffuse in both cells, or septum. The differences in localization of MreB_6xH between treatment conditions at each stage of division were statistically analyzed using a chi-squared test to reveal that the changes resulting from drug treatments were statistically significant when compared to UTD (* = p<0.0001). Pretreatment of cells with diNN and A22 resulted in a statistically significant difference in MreB localization in pre-division intermediates (round cells) when compared to pretreatment with diNN alone (** = p<0.02). For (A) and (B), data were pooled from two independent experiments. (C) Representative images of the different division intermediates from untreated cells or from cells treated with drugs as indicated in the figure. Scalebar = 2 μm

Evaluating the localization of the ectopically expressed protein under these conditions, we observed that MreB_6xH was primarily in a polar cluster in pre-division “round” intermediates (Figs. 7B&C). Although MreB_6xH exhibited a septal localization profile in the majority of early division intermediates, a subset of these intermediates retained MreB_6xH at the leading edge of the daughter cell suggesting there is a slow transition of MreB from the leading edge of the daughter to the septum at this stage of division (Figs. 7B&C). In late budding intermediates, MreB_6xH was primarily septal, and it was mostly diffuse at the two-cell stage (Figs. 7B&C). A very similar array of MreB localization profiles was observed when we induced the expression of MreB_6xH in infected cells at 21hpi and partially purified *Chlamydia* at 22hpi and determined the distribution of the fusion (Supp. Fig. 6). These results suggest that MreB, like Cls, transitions between different cellular compartments during the polarized budding process of *Chlamydia*.

To assess the effect of diNN on the localization profile of MreB, partially purified *Chlamydia* were pre-treated with 5μM diNN for 30 minutes in axenic media then induced with aTc for 1 hour in the continued presence of the drug. This drug treatment resulted in a diffuse pattern of localization of MreB_6xH in the majority of round, early, and mid/late division intermediates (Figs. 7B&C). Quantification of these analyses revealed a statistically significant effect of diNN on the distribution of MreB_6xH at each stage of division compared to the untreated control (Fig. 7B). These results indicate that the localized assembly of MreB in dividing chlamydiae is dependent upon the maintenance of CL-rich membrane microdomains.

It was possible that a preexisting pool of endogenous polymeric MreB serves as a nucleation site for the assembly of newly synthesized MreB_6xH in a subset of the dividing cells treated with diNN. Therefore, in addition to diNN, we also pretreated cells with A22, to depolymerize this preexisting pool of endogenous MreB. The A22 was then removed and MreB_6xH was induced for 1 hour in the continued presence of diNN. Although there was a reduction in the number of cells in the population undergoing division when they were treated in this way (Fig. 7A), the results observed in those cells undergoing division were similar to the treatment with diNN alone (Figs. 7A-C). However, pretreatment with A22 resulted in a further significant reduction in the number of round (pre-division) intermediates with polar MreB_6xH, suggesting that, at least at this stage of division, a preexisting pool of polymeric MreB may serve as a nucleation site for the assembly of newly synthesized MreB in the presence of diNN. The observation that the A22 pretreatment did not have any significant effect beyond the effect of diNN alone on the assembly of newly synthesized MreB at the septum (yellow in bar graph of Fig. 7B) at any stage of division was unexpected. To determine whether a longer treatment with A22 would eliminate the septal assembly of MreB, we pretreated cells with A22 for 30 minutes, then MreB_6xH was induced for 1 hour in the continued presence of A22. However, this prolonged treatment with A22 resulted in a further reduction in the number of *Chlamydia* in the population undergoing division (data not shown) making it impossible to assess the effect of A22 on the assembly of MreB at later stages of division. While we were unable to fully define the mechanisms that regulate the assembly of septal MreB, our results indicate a critical role for CL-rich membrane domains in directing the localized assembly of MreB in round, early, and mid/late division intermediates. Furthermore, the formation of these CL-rich membrane microdomains depends upon the polarized synthesis of CL by Cls.

## Discussion

In model organisms like *E. coli* or *B. subtilis* that use a FtsZ-dependent binary fission mechanism to divide, there are conserved systems to ensure proper placement of the FtsZ ring in the middle of the cell. These include, for example, the MinCDE and nucleoid occlusion (Noc) systems [43–46]. *Chlamydia* has no annotated or otherwise obvious orthologs to these proteins (ParA, a chromosome partitioning ATPase that *Chlamydia* encodes, shares homology with MinD) [10]. *Chlamydia* is unique amongst characterized bacteria in using an MreB-dependent polarized cell division process [11, 16]. Although much recent work has identified the predicted components of the chlamydial divisome [11,12,14,31], it is unclear how a division site is selected given that *Chlamydia* are coccoid bacteria with apparent uniformity in their negative membrane curvature. An additional issue for *Chlamydia* is that MreB is typically excluded from areas of negative membrane curvature, thus how it is recruited to the inner leaflet of the inner membrane at a distinct site was unclear prior to this study.

Our data indicate that CL-rich membrane domains are necessary for the localized assembly of MreB in pre-(round), early, and mid/late division intermediates. We hypothesize that this polar population of CL either directly recruits MreB or drives localized membrane deformation to enable MreB recruitment to the site where the daughter cell will bud (modeled in Figure 8). CL has a small head group with 4 acyl chains that allows it to be packed into areas of high curvature [24]. Although seemingly counterintuitive, localized deposition of CL in the inner leaflet would induce membrane bending that necessarily creates areas of *positive* curvature in the paired outer leaflet of the membrane and in the areas directly flanking the inner leaflet where CL was deposited. Such areas would favor MreB recruitment and subsequent peptidoglycan synthesis to initiate the division process. Continued outgrowth of the budding daughter cell would, however, necessitate CL in the outer leaflet at the “junction” where the daughter cell arises from the mother cell since this area will also have curvature constraints.

**Figure 8.**
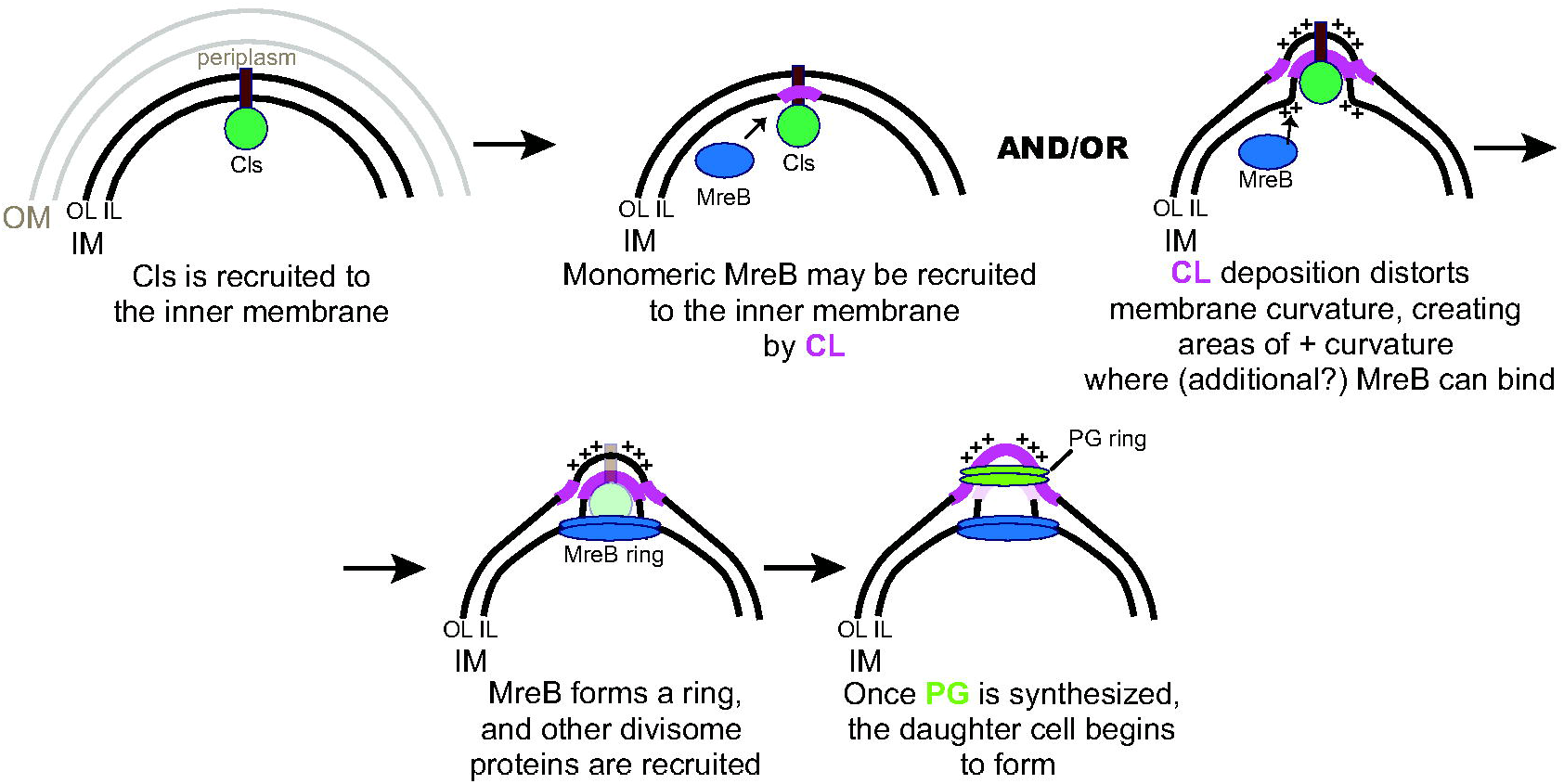
Model for the function of cardiolipin (CL) and Cls in chlamydial cell division. How Cls is recruited to the membrane at a distinct site is not understood but requires the transmembrane domain. An initial deposition of CL may either directly recruit MreB or induce membrane bending that recruits MreB, and these are not mutually exclusive. Once MreB forms a ring, it recruits subsequent cell division machinery to allow peptidoglycan (PG) synthesis in the periplasm. OM = outer membrane IM = inner membrane OL = outer leaflet of IM IL = inner leaflet of IM ++ = areas of positive membrane curvature in each leaflet of the IM.

To test how a polarized distribution of CL could arise in *Chlamydia*, we assessed various parameters associated with localization of the chlamydial CL synthase. At the onset of division, Cls exhibited a polarized pattern of localization at the side of the cell where the daughter cell forms. Cls remained associated with the leading edge of the daughter cell at later stages of the division process in support of our model. The restricted localization of Cls overlapped the distribution of NAO-labeled aPLs, and Cls and NAO-labeled aPLs were redistributed to the mother cell by diNN, a drug that disrupts CL-rich membrane microdomains. Importantly, the localization of a neutral phospholipid (phosphatidylethanolamine; PE) synthase, PsdD, was restricted to the mother cell membrane distal to the division site, and its localization was unaffected by diNN. These data suggest that general PL membrane synthesis may be restricted to the mother cell during the division process, which is consistent with the function of PE as a major cell membrane constituent in *Chlamydia* [25, 26]. Conversely, the deposition of CL is restricted to a polarized site in *Chlamydia* associated with the dividing (daughter) side of the cell. It remains unclear at this time if the recruitment of Cls to a polarized site is regulated or if its insertion into the membrane creates spontaneous symmetry breaking that, in turn, locks in the polarization of the organism. Given the polarity observed in the EB form [16], it is likely the former, but our data cannot currently discriminate between these possibilities.

Our model predicts that Cls acts epistatically to MreB and other divisome machinery in the division process. Our data support such a model. Restricted localization of Cls_6xH either before or after treating infected cells with the MreB or PG-disrupting drugs, A22 or D-cycloserine (D-CS), respectively, was not altered. Conversely, MreB membrane localization was disrupted by A22, as previously noted [13], with a concomitant loss of PG signal. D-CS, as expected [42], did not impact MreB but did disrupt PG ring formation, resulting in aberrantly enlarged organisms. Overall, these data suggest that Cls is epistatic to MreB during cell division. Moreover, our inability to detect a physical interaction between Cls and divisome components by two hybrid assays suggests that it is the activity of Cls (i.e. generating CL) and not its ability to recruit specific protein factors that is important for its function in any cell division context. Importantly, our results with diNN-treated *Chlamydia* support our proposed model and suggest that the establishment of CL-rich membrane microdomains is critical for the localized assembly of MreB during chlamydial division.

It is not clear on which side of the inner membrane CL exerts its function as PLs can be flipped across membranes [47]. Importantly, our model posits a function for CL on *both* leaflets depending on the stage of daughter cell outgrowth. Determining lipid localization, particularly to a single leaflet is difficult, and the small size of chlamydial RBs preclude using light-based microscopy techniques to address this question. To investigate the distribution of CL in the chlamydial membrane, we expressed a characterized mitochondrial CL binding protein, C11orf83 [29], within different compartments of *Chlamydia*. Of note, C11orf83 also binds phosphatidic acid (PA) [29], which is present at low levels as a short-lived PL precursor in bacteria that is incorporated into CDP-diacylglycerol in the PL synthesis pathway [36, 37]. The full-length C11orf83 exhibited a diffuse localization profile in *Chlamydia* and had no obvious impact on chlamydial growth or morphology. In contrast, by tethering the full-length C11orf83 to the inner leaflet of the inner membrane using the Cls TM domain, there was a dramatic and negative impact on *Chlamydia*. However, given the possibility of C11orf83’s binding PA at the inner leaflet, we cannot with absolute certainty assign the negative impact on *Chlamydia* to disruption of CL. Strikingly, when C11orf83 was tethered to the outer leaflet of the inner membrane within the periplasm, this severely disrupted chlamydial growth and morphology as well. We are unaware of any reports demonstrating free PA in the outer leaflet of the inner membrane or the periplasm more generally in bacteria, thus it is likely that the negative impacts of C11orf83 in the periplasm are due to its binding of CL. This was not a general effect associated with overexpressing a protein in the periplasm as we did not observe such effects using mCherry tethered to the outer leaflet. Collectively, these data suggest a critical function for CL in both leaflets of the inner membrane and support our model.

Other published data related to divisome components and PG are consistent with our proposed model that an initial deposition of MreB and PG, driven by localized CL synthesis in the membrane, occurs before MreB or PG rings are observed. For example, we show here that MreB forms a polarized cluster at the MOMP-enriched side of the cell prior to division and a ring at the base of the daughter cell as it grows. More recently, we demonstrated specific functions for the PG transpeptidases PBP2 and PBP3 in daughter cell growth, showing that inhibition of PBP2 prevents the initiation of daughter cell formation with only a patch of polarized PG detectable underneath the MOMP label in treated organisms [17]. MreB is epistatic to PBP2 and PBP3 [11].

How polarity is established in *Chlamydia* is an interesting question. In other organisms, specific protein factors like bactofilins have been associated with polar localization. Interestingly, *Chlamydia* has a bactofilin ortholog, but, as we recently characterized, it is not associated with division but rather with cell size [48]. Therefore, further work is required to understand how Cls is recruited to a discrete site and whether specific protein factors or other context-dependent cues drive its localization. Other questions under investigation in our labs include how membrane synthesis changes as the daughter cell grows and how these changes are resolved at the completion of the division process. Given the unique biology of *Chlamydia*, it is possible that they have once again co-opted conserved systems for their specific needs.

## Materials and Methods

### Organisms and Cell Culture

HeLa (ATCC, Manassas, VA), HEp2, and McCoy (kind gift of Dr. Harlan Caldwell) cells were cultured in Dulbecco’s Modified Eagle Medium (DMEM; Invitrogen, Waltham, MA) containing 10% fetal bovine serum (FBS; Hyclone, Logan, UT) and 10 μ MA) at 37 ℃ with 5% CO_2_. Wild-type *C. trachomatis* serovar L2/434/Bu was cultured in HeLa cells, and EBs were purified from cell lysates through a renografin density gradient as described elsewhere [49]. *Chlamydia trachomatis* serovar L2 lacking the endogenous plasmid (-pL2) (kind gift of Dr. Ian Clarke) was infected and propagated in McCoy cells for use in transformations. HeLa cells were infected with chlamydial transformants in DMEM containing 10% FBS, 10 μ g/mL cycloheximide. All cell cultures and chlamydial stocks were routinely tested for Mycoplasma contamination using the Mycoplasma PCR detection kit (Sigma, St. Louis, MO). All chemicals and antibiotics were obtained from Sigma unless otherwise noted.

### Cloning

The plasmids and primers used in this study are described in Supplemental Table 1. The wild-type or mutant chlamydial *cls, psdD*, and *mreB* genes were amplified by PCR with Phusion DNA polymerase (NEB, Ipswich, MA) using 10 ng *C. trachomatis* L2 genomic DNA as a template. Some gene segments were directly synthesized as a gBlock fragment (Integrated DNA Technologies, Coralville, IA). The PCR products were purified using a PCR purification kit (Qiagen, Hilden, Germany). The HiFi Assembly reaction master mix (NEB) was used according to the manufacturer’s instructions in conjunction with plasmids pBOMB4-Tet (kind gift of Dr. Ted Hackstadt [50]) cut with EagI and KpnI or the BACTH vector pSTM25 digested with KpnI and BamHI [38]. All plasmids were dephosphorylated with alkaline phosphatase (FastAP; ThermoFisher) prior to use in the HiFi reaction. The products of the HiFi reaction were transformed into NEB-10β (chlamydial transformation) or NEB-5αI (BACTH) competent cells (NEB), plated on appropriate antibiotics, and plasmids were subsequently isolated from individual colonies grown overnight in LB broth by using a mini-prep kit (Qiagen). All plasmids were verified for correct size by digest, and inserts were sequenced.

### Bioinformatics Analysis

Sequences for *Chlamydia trachomatis* serovar L2/434, different chlamydial species, and *Bacillus subtilis* were obtained from the NCBI database (https://www.ncbi.nlm.nih.gov/) and for *E. coli* MG1655 from Ecocyc database (https://ecocyc.org/) [51]. Protein sequence alignment was performed using Clustal Omega website (https://www.ebi.ac.uk/Tools/msa/clustalo/) [52] and the ESPript3 program (http://espript.ibcp.fr) [53]. TOPCONS [54] and TMHMM [55] were used for transmembrane domain prediction.

### Nucleic acid extraction and RT-qPCR

Total RNA and DNA were extracted from *C. trachomatis* L2/434Bu-infected HEp2 cells plated in 6-well dishes as described previously [56]. Briefly, for RNA, cells were rinsed one time with PBS, then lysed with 1mL Trizol (Invitrogen) per well. Total RNA was extracted from the aqueous layer after mixing with 200μL per sample of chloroform following the manufacturer’s guidelines. Total RNA was precipitated with isopropanol. Purified RNA was rigorously DNased using DNAfree (Ambion) according to the manufacturer’s guidelines prior to synthesis of cDNA using SuperScript III (Invitrogen) following the manufacturer’s guidelines. For DNA, cells were rinsed one time with PBS, trypsinized, and pelleted before resuspending each pellet in 500μL PBS. Each sample was split in half (i.e. 250μL), and genomic DNA was isolated from each duplicate sample using the DNeasy extraction kit (Qiagen) according to the manufacturer’s guidelines.

Quantitative PCR to detect genomic DNA (gDNA) levels of *C. trachomatis* and RT-qPCR to detect the indicated transcripts were performed as described previously using SYBR Green [56]. For gDNA levels, 150ng of each sample was used in 25μL reactions using standard amplification cycles on a Quantstudio3 thermal cycler (Applied Biosystems) followed by a melting curve analysis. For cDNA levels, equal volumes of each cDNA reaction were used in 25μL reactions under standard amplification cycle conditions with melting curve analysis. Transcript levels were normalized to genomes and expressed as ng cDNA/gDNA.

### Transformation of Chlamydia trachomatis

McCoy cells were plated in a six-well plate at a density of 1 x 10^6^ cells per well the day before beginning the transformation procedure. *C. trachomatis* serovar L2 lacking its endogenous plasmid (-pL2) was incubated with 2 μg plasmid in Tris-CaCl_2_ buffer (10 mM Tris-Cl pH 7.5, 50 mM CaCl_2_) for 30 min at room temperature [6, 8]. During this step, the McCoy cells were washed with 2 mL 1X Hank’s Balanced Salt Solution (HBSS) media containing Ca^2+^ and Mg^2+^ (Gibco). After that, McCoy cells were infected with the transformants in 2 mL HBSS per well. The plate was centrifuged at 400 x g for 15 min at room temperature and incubated at 37℃ for 15 min. The inoculum was aspirated, and DMEM containing 10% FBS and 10 μg/mL gentamicin was added per well. At 8 h post infection (hpi), the media was changed to media g/mL cycloheximide and 1 or 2 U/mL penicillin G, and the plate was incubated at 37 ℃ until 48 hpi. At 48 hpi, the transformants were harvested and used to infect a new McCoy cell monolayer. These harvest and infection steps were repeated every 48 hpi until mature inclusions were observed. DNA was isolated from transformants using a Genomic DNA isolation kit (DNeasy; Qiagen) and used to transform NEB-10β competent cells to reverify the plasmid as described above for size and insert sequence fidelity.

### Indirect Immunofluorescence (IFA) Microscopy

HeLa cells were seeded in 24-well plates on coverslips at a density of 1.5 x 10^5^ cells per well the day before infection. Chlamydial strains expressing wild-type Cls or MreB with a six-histidine tag at the C-terminus were used to infect HeLa cells in DMEM media containing 1 U/ml penicillin G and 1μ g/mL cycloheximide. Anhydrotetracycline (aTc) was added at the indicated concentration at the indicated time. The coverslips of infected cells were washed with 1X DPBS and fixed with fixing solution (3.2% formaldehyde and 0.022% glutaraldehyde in 1X DPBS [16]) for 2 min at 10.5 hpi or 20 hpi. To observe the effect of A22 and D-cycloserine (DCS), we added 75 μg/mL DCS at 18 hpi or 4 hpi, respectively. The samples were then washed three times with 1X DPBS and permeabilized with ice-cold 90% methanol for 1 min. Afterwards, the fixed cells were labeled with primary antibodies including goat anti-major outer-membrane protein (MOMP; Meridian, Memphis, TN), rabbit anti-chlamydial MreB antibody (custom anti-peptide antibody directed against the C-terminus of *C. trachomatis* serovar L2 MreB; ThermoFisher), or rabbit anti-six histidine tag (Genscript, Nanjing, China and Abcam, Cambridge, MA, respectively) as indicated. Donkey anti-goat (647 or 488) or donkey anti-rabbit (594) secondary antibodies (Invitrogen) were used to visualize the primary antibodies.

Coverslips were observed by using either a Nikon Ti2 spinning disc confocal microscope using a 60X lens objective or a Zeiss AxioImager.Z2 equipped with an Apotome2 using a 100X lens objective.

### Peptidoglycan (PG) staining

PG was labelled with D-amino acid dipeptide probes and click chemistry as previously described [42]. Briefly, HeLa cells were infected with *C. trachomatis* containing an anhydrotetracycline (aTc)-inducible vector encoding chlamydial Cls_6xH or MreB_6xH. 1 mM EDA-DA, which is the D-amino acid dipeptide probe, was added at 0 hpi. The constructs were induced with 10 nM anhydrotetracycline (aTc) at 16 hpi. At 20 hpi, cells were washed 3 times with 1X DPBS and fixed with 100% methanol for 5 min. After washing the samples three times with 1X DPBS, the cells were permeabilized with 0.5% Triton X-100 for 5 min. The samples were blocked with 3% Bovine Serum Albumin (BSA) for 1 hour. The PG was labeled by using the Click-iT cell reaction buffer kit according to the manufacturer’s instructions (Invitrogen, Waltham, MA).

### Inclusion forming unit assays

HeLa cells were infected with C*. trachomatis* serovar L2 transformed with a plasmid encoding an aTc-inducible gene as indicated. At 8 hpi, aTc was added to the culture media at the indicated concentration. At 24 hpi the HeLa cells were dislodged from the culture dishes by scraping and collected into microcentrifuge tubes. Suspensions were centrifuged at 4°C for 30 min, the supernatant was removed by aspiration, and the pellet was resuspended in 1mL 2 sucrose-phosphate (2SP) solution [6] and frozen at -80°C. At time of secondary infection, the chlamydiae were thawed on ice and vortexed. Cell debris was pelleted by centrifugation for 5 min at 1k xg, 4°C. The *C. trachomatis* elementary bodies (EBs) in the resulting suspension were serially diluted and used to infect a monolayer of HeLa cells in a 24-well plate. The secondary infection was allowed to grow at 37°C with 5% CO_2_ for 24 h before they were fixed, labeled for immunofluorescence microscopy with goat anti-MOMP antibody and a secondary donkey anti-goat antibody labeled with Alexa Fluor 594, and counted. Titers were enumerated by calculating the total number of inclusions per field based on counts from 20 fields of view. Three independent replicates were performed, and the totals for each experiment were averaged. Results were normalized as a percentage of the uninduced samples for each time point. Student’s two-tailed t test to compare the induced samples to the uninduced samples was performed using the averages of each biological replicate.

### Localization of Cls_6xH and anionic phospholipids in partially purified *C. trachomatis*

HeLa cells were infected with C*. trachomatis* serovar L2 transformed with a plasmid encoding an aTc-inducible Cls_6xH. At 20hpi, 10nM aTc was added to the culture media, and the HeLa cells were then harvested from the culture dishes by scraping at 22hpi. The HeLa cells were pelleted, and *Chlamydia* were released from infected cells as described previously [6]. Briefly, infected HeLa cells were resuspended in 1mL of 0.1x PBS and vortexed in tubes containing 0.1mM glass beads. The lysate was centrifuged at 1000rpm for 3 minutes and 20μL of the supernatant was transferred to a glass slide and mixed with an equal volume of 2x fixative (6.4% formaldehyde, 0.044% glutaraldehyde in PBS) and incubated for 10 min at room temperature. The cells were washed 3x in PBS and permeabilized by incubation with PBS containing 0.1% TX-100 for 1 minute. Following washing in PBS, the cells were incubated with goat polyclonal antibodies directed against MOMP and rabbit polyclonal antibodies directed against 6xHis tag (Abcam) for 1 hour. The cells were washed and incubated with donkey anti-goat IgG conjugated to Alexa 594 and donkey anti-rabbit IgG conjugated to Alexa 488. Cells were again washed and imaged using Zeiss AxioImager2 microscope equipped with a 100x oil immersion PlanApochromat lens. Images were deconvolved using the deconvolution algorithm in the Zeiss

Axiovision 4.7 software. Alternatively, aTc-induced cells labeled with MOMP antibodies were incubated with 250nM NAO (Invitrogen) to visualize the distribution of anionic phospholipids following Cls_6xH induction. Virtually identical results were obtained when infected HeLa cells were harvested, resuspended in 1mL of 1x PBS, and mixed with an equal volume of 2x fixative and incubated for 10 min at room temperature. Fixed cells were washed 2x in PBS then resuspended in 1 mL of PBS and vortexed with glass beads. The lysate was centrifuged at 1000rpm for 3 minutes and 20μL of the supernatant was transferred to a glass slide and stained as described above (data not shown). In some instances, the distribution of Cls_6xH and NAO staining were visualized in the same cell that was also stained with HOECHST 33342 to visualize DNA.

Z-stacks were collected during image acquisition that extended above and below the dividing cell. The largest diameter of the nascent daughter cell and the progenitor mother cell was determined using the measuring tool in the Zeiss AxioVision 4.7 software. This value was used to estimate the volume of the daughter and mother cell according to the formula, v=4/3πr^3^. For all experiments we defined several intermediates in the polarized division process. Pre-division intermediates (round cells) had no obvious outgrowth of the budding daughter cell, in early division intermediates the daughter cell volume was <15% of the mother cell volume, in mid/late division intermediates the daughter cell volume was between 15-80% of the mother cell volume, and in the two-cell stage the daughter cell volume was >80% of the mother cell volume.

Effect of the amphiphilic aminoglycoside, 3’,6-dinonylneamine (diNN), on the localization of Cls_6xH and anionic phospholipids in *C. trachomatis* grown in axenic media

HeLa cells were infected with C*. trachomatis* serovar L2 transformed with a plasmid encoding an aTc-inducible Cls_6xH. At 22hpi, the HeLa cells were harvested as described above, and cells were lysed by vortexing in 1mL of 0.1x PBS in tubes containing 0.1mM glass beads. The lysate was centrifuged at 1000rpm for 3 minutes and 20 μL of the supernatant was transferred to a glass slide where it was mixed with 100μL of axenic media composed of Opti-MEM^TM^ (Gibco) supplemented with 1mM L-Glutamine, 1mM glucose-6-phosphate (Moltox), and 10nM aTc. The slide was transferred to a humidified chamber and incubated at 37 °C for 1.5 hours in a CO_2_ incubator. The cells were then fixed by mixing the cells with an equal volume of 2x fixative and incubating for 10 min at room temperature. Fixed cells were permeabilized by incubating in PBS containing 0.1% TX-100 for 1 min, and the localization of Cls_6xH and anionic phospholipids in aTc-induced cells was determined as described above. To determine the effect of the amphiphilic aminoglycoside, diNN, on the localization of Cls_6xH and anionic phospholipids, 5μM diNN was present in the axenic media throughout the aTc induction.

### Effect of 3’,6-dinonylneamine (diNN) and A22 on the localization of MreB_6xH in *C. trachomatis* grown in axenic media

HeLa cells were infected with C*. trachomatis* serovar L2 transformed with a plasmid encoding an aTc-inducible MreB_6xH. At 22hpi, *Chlamydia* were partially purified as described above and the cells were incubated in axenic media containing 10nM aTc for 1 hour at 37°C in a CO_2_ incubator. The cells were then fixed and the distribution of MreB_6xH in dividing cells was assessed by staining with MOMP and 6xHis antibodies. The effect of 5μM diNN and 75μM A22 on MreB_6xH localization was assessed by pretreating cells with diNN alone or diNN + A22 for 30 minutes prior to a one hour induction with aTc in the presence of diNN alone.

### Quantification of Cls_6xH, MreB_6xH, and NAO staining profiles in *C. trachomatis* induced with aTc during infection or in axenic media

*C. trachomatis* in which Cls_6xH or MreB-6xH was induced were stained with MOMP antibodies and the 6xHis-tag antibody or NAO. For each condition, the Cls_6xH, MreB_6xH, and NAO staining profiles were imaged in at least 50 cells for each stage of division. For each condition, the MreB_6xH localization profile was imaged in at least 100 cells for each stage of division. For some of the drug treatments fewer than 100 cells were counted because certain division intermediates were present at very low levels following drug treatment. Localization profiles for the Cls_6xH, MreB_6xH, and NAO were categorized and the effect of the drugs on the localization profiles determined and statistically analyzed using a chi-squared test.

## Acknowledgments

The authors would like to acknowledge Dr. H. Caldwell (NIH) for cell lines, Dr. I. Clarke (Univ. Southampton) for the -pL2 strain of *C. trachomatis*, and Dr. Ted Hackstadt (RML/NIH) for the aTc-inducible plasmid. The authors are grateful to Dr. Jean-Luc Décout (Université Grenoble Alpes) for sharing the diNN compound. SPO and JVC were supported by awards from the National Science Foundation (1817583 to SPO and 1817578 to JVC) and the National Institutes of Health/NIGMS (1R35GM124798-01 to SPO).

This publication’s contents are the sole responsibility of the authors and do not necessarily represent the official views of the NSF, NIH, or NIGMS.

## Supplemental Material

**Supplemental Table S1.** List of primers and plasmids used in the current study.

**Supplemental Figure S1.** (A) Alignment of *ct284/cls* orthologs from multiple pathogenic chlamydial species. The numbering is for the *C. trachomatis* protein. (B) Transcriptional analysis of *cls* during the *C. trachomatis* L2 developmental cycle. Cells were infected with wild-type *C. trachomatis* L2/434Bu, and total RNA and DNA were collected at the indicated time points. Transcript levels were normalized to genomic DNA levels and are expressed as ng cDNA/gDNA. Shown are data from one of two experiments.

**Supplemental Figure S2.** Effect of extended overexpression of the Cls_TM_GFP construct on chlamydial morphology. Cells were infected as described in the legend of Figure 3, and expression of Cls_TM_GFP was induced or not at 8hpi with 2 or 10nM aTc. Cells were fixed at 24hpi and processed for immunofluorescence using antibodies against the major outer membrane protein (MOMP). DNA was visualized with DAPI. Images were acquired on a Zeiss AxioImager.Z2 equipped with an Apotome2 using a 100X lens objective. Images are representative of at least three independent experiments. Scalebar = 2 μm

**Supplemental Figure S3.** Effect of diNN on localization of Cls_TM_GFP. Samples were collected and processed as described in the legend to Figure 4. (A) Representative images of diNN-treated Cls_TM_GFP expressing bacteria at each stage of division. (B) 50 individual cells from Early, Mid/late and Two-cell stages of division from untreated and diNN-treated cultures were assessed for the localization of Cls_TM_GFP. Localization profiles were categorized into leading edge of the budding daughter cell, diffuse in mother cell, diffuse in one cell, diffuse in both cells, or discrete in one cell. The differences in Cls_TM_GFP localization between treatment conditions at each stage of division were statistically analyzed using a chi-squared test to reveal that the changes observed during diNN treatment were statistically significant (p<0.001).

**Supplemental Figure S4.** Effect of diNN on localization of PsdD_6xH. Samples were collected and processed as described in the legend to Figure 4. Representative images of untreated (UTD) or diNN-treated PsdD_6xH expressing bacteria at each stage of division are shown. 50 individual cells from each stage of division from untreated and 35 cells from diNN-treated cultures were assessed for their localization to the leading edge of the budding daughter cell, the mother cell, or both cells for Early and Mid/Late stages, whereas, for the Two-cell stage, discrete localization in one cell or diffuse localization in one or both cells was quantified. The differences in localization of PsdD_6xH between treatment conditions at each stage of division were statistically analyzed using a chi-squared test to reveal that the changes observed during diNN treatment were not statistically significant.

**Supplemental Figure S5.** Cls localization is unaffected by disrupting MreB activity. Cells were either pre-treated with A22 for 2h prior (Pre) to inducing (A) Cls_6xH or (B) MreB_6xH expression at 16hpi or treated after inducing (Post) at 18hpi. See legend to Figure 6 for more details.

**Supplemental Figure S6.** MreB_6xH localization in RBs isolated from infected cultures. As described in the legend of Figure 2B, cells were infected with the aTc-inducible MreB_6xH transformant, and expression of the construct was induced prior to isolating and partially purifying RBs from infected cells. The localization of MreB_6xH in the division intermediates is shown. See also Figure 7B to compare the localization of MreB_6xH induced after purifying RBs. Scalebar = 2 μm

## Notes

### Competing Interest Statement

The authors have declared no competing interest.

