## Supplementary figures and images for "Localized Cardiolipin Synthesis is Required for the Assembly of MreB During the Polarized Cell Division of *Chlamydia trachomatis*"

### Supplemental Figure 1

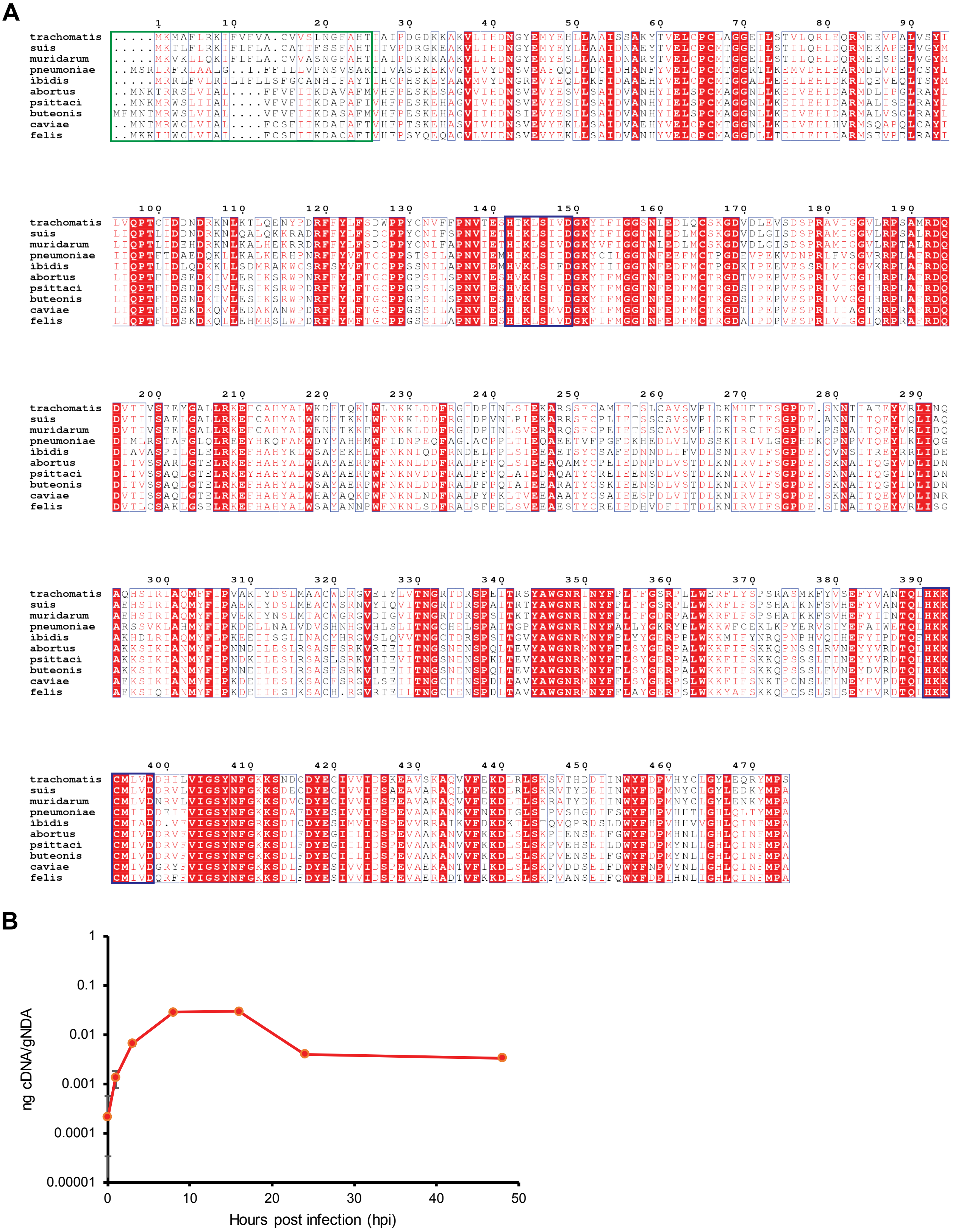


**Supplemental Figure 1**

### Supplemental Figure 2

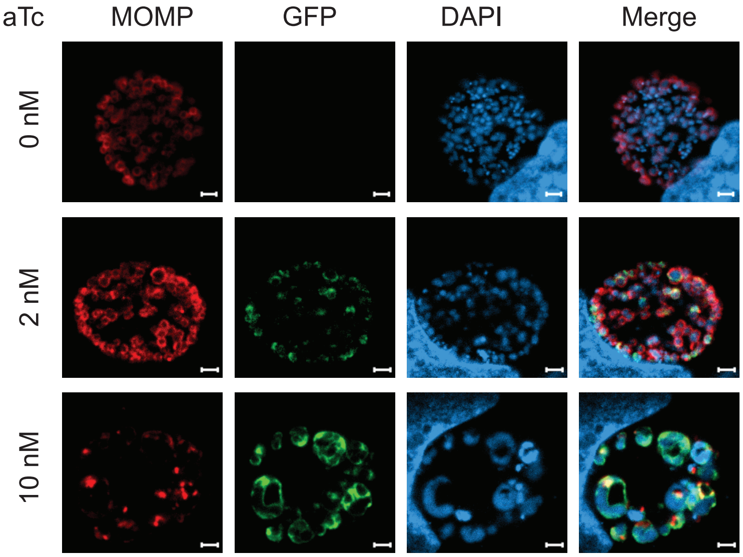


**Supplemental Figure 2**

### Supplemental Figure 3

**A**

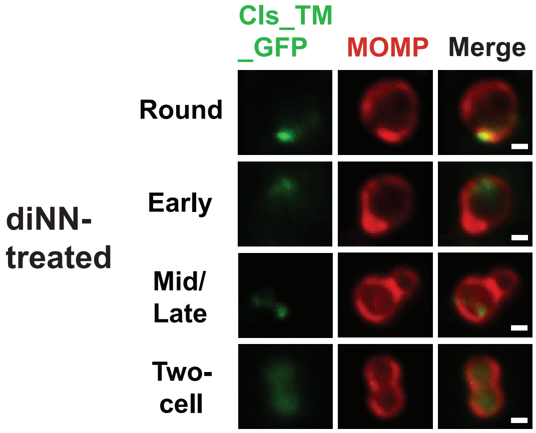


**B**

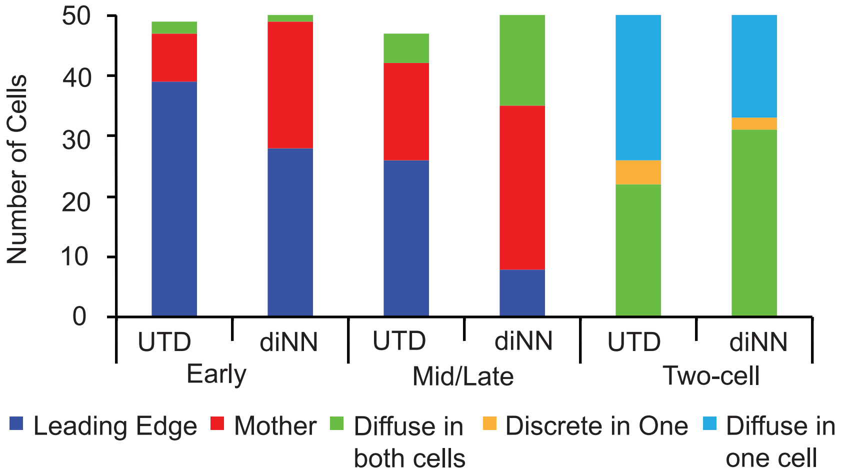


**Supplemental Figure 3**

### Supplemental Figure 4

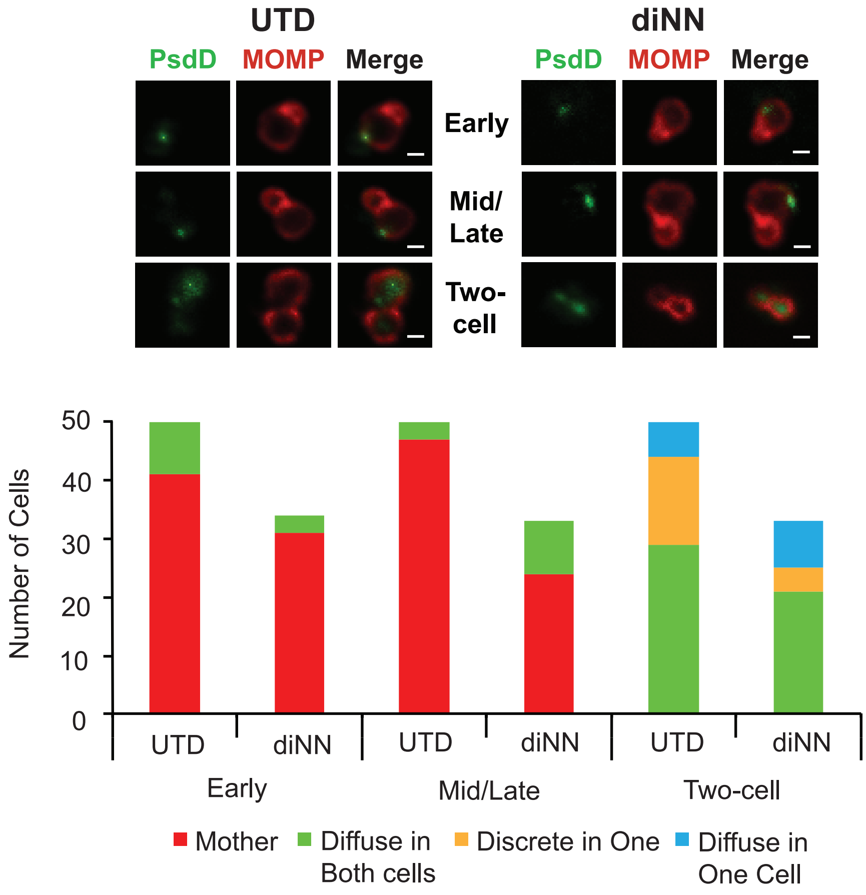


**Supplemental Figure 4**

### Supplemental Figure 5

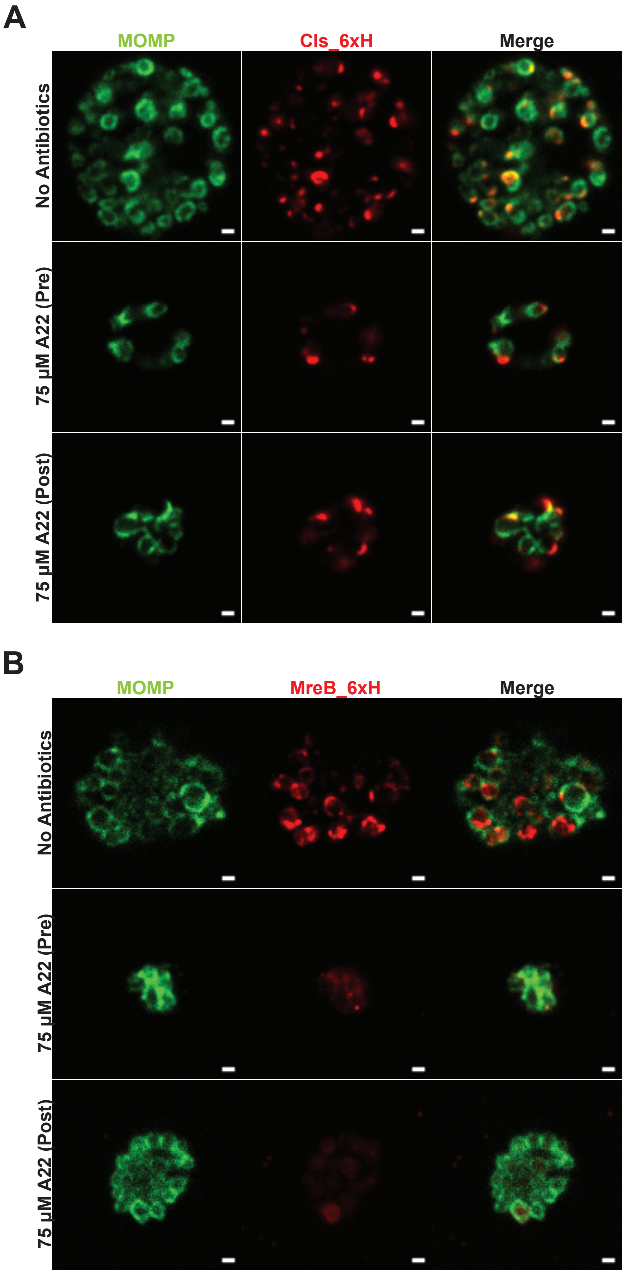


**Supplemental Figure 5**

### Supplemental Figure 6

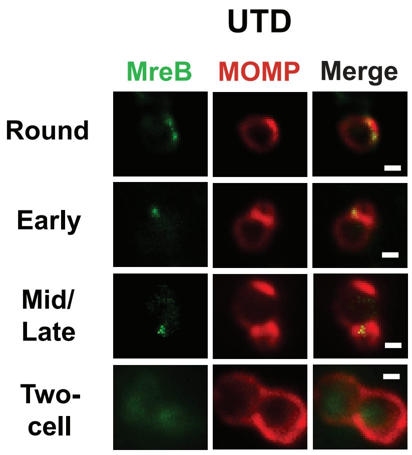


Supplemental Figure 6
