## Supplemental Table 1 for "Localized Cardiolipin Synthesis is Required for the Assembly of MreB During the Polarized Cell Division of *Chlamydia trachomatis*"

**Supplementary table 1. List of Plasmids and Primers**

| Plasmid | Relevant genotype | Ori | Source or Reference |
|---|---|---|---|
| pBOMB4-Tet::L2 | *bla* P*tet*::*mCherry* $P_{Nm}$::*gfp* | pUC19 | 1 |
| pBOMB-cls_6xH::L2 | *bla* P*tet*::*cls_6xH* $P_{Nm}$::*gfp* | pUC19 | This study |
| pBOMB-ΔN25_cls_6xH::L2 | *bla* P*tet*::ΔN25_*cls_6xH* $P_{Nm}$::*gfp* | pUC19 | This study |
| pBOMB-psdD_6xH::L2 | *bla* P*tet*::*psdD_6xH* $P_{Nm}$::*gfp* | pUC19 | This study |
| pTLR2-*mreB*_6xH | *bla* P*tet*::*mreB*_6xH $P_{dnaKmut}$::*mKate2* | ColE1 | 2 |
| pBOMBmC::L2 | *bla* P*tet*::*mCherry* $P_{Nm}$::*mCherry* | pUC19 | 3 |
| pBOMBmC-cls_TM_GFP::L2 | *bla* P*tet*::*cls_TM_GFP* $P_{Nm}$::*mCherry* | pUC19 | This study |
| pBOMB-c11orf83_6xH::L2 | *bla* P*tet*::*c11orf83_6xH* $P_{Nm}$::*gfp* | pUC19 | This study |
| pBOMB-cls_TM_c11orf83_6xH::L2 | *bla* P*tet*::*cls_TM_c11orf83_6xH* $P_{Nm}$::*gfp* | pUC19 | This study |
| pBOMB-TM_c11orf83_6xH::L2 | *bla* P*tet*:: $TM_{Ec\_OppB}$*_c11orf83_6xH* $P_{Nm}$::*gfp* | pUC19 | This study |
| pBOMB-TM_mCherry::L2 | *bla* P*tet*:: $TM_{Ec\_OppB}$*_mCherry* $P_{Nm}$::*gfp* | pUC19 | This study |
| pSTM25 | *aadA* P*lac*:: *t25*-$TM_{Ec\_OppB}$ | p15A | 4 |
| pSTM25-cls | *aadA* P*lac*:: *t25*-$TM_{Ec\_OppB}$-*cls* | p15A | This study |

| Primer name | Sequence | Features | Usage |
|---|---|---|---|
| cls/(pBOMB)/5' | gatctaaagaggagaaaggatctgcATGAAAATGGCTTTTTACG | lower case for plasmid overlap construction | For amplification of cls into pBOMB |
| cls_6xH/(pBOMB)/3' | tttgaatggtcgaccggtacctgcattaatggtgatggtgatggtgAGATGGCATGTATCTCTGTTC | lower case for plasmid overlap construction; adds 6xH sequence to cls | For amplification of cls into pBOMB |
| psdD/(pBOMB)/5' | gatctaaagaggagaaaggatctgcATGGCAGCGCGGGAAATG | lower case for plasmid overlap construction | For amplification of psdD into pBOMB |
| psdD_6xH/(pBOMB)/3' | tttgaatggtcgaccggtacctgcattaatggtgatggtgatggtgTGAAGAGAAACGTTTTCCTAACGATTG | lower case for plasmid overlap construction; adds 6xH sequence to cls | For amplification of psdD into pBOMB |
| cls/(pSTM25)/5’ | acgcacaagggcctctagagATAGCTATTCCGGATGGAG | lower case for plasmid overlap construction | For amplification of cls into pSTM25 |

| | | | |
|---|---|---|---|
| cls/(pSTM25)/3’ | attcttagttacttaggtacttaAGATGGCATGTATCTC | lower case for plasmid overlap construction | For amplification of cls into pSTM25 |
| N26-cls/(pBOMB)/5’ | aaagaggagaaaggatctgcATGGCTATTCCGGATGGAGAC | lower case for plasmid overlap construction | For amplification of N26_cls into pBOMB |
| cls_TM/(pBOMBmC)/5’ | aaagaggagaaaggatctgcATGAAAATGGCTTTTTTACGG | lower case for plasmid overlap construction | For amplification of cls_TM fragment |
| cls_TM/(GFP)/3’ | ctcctttactAGTGTGCGCAAAACCATTC | lower case for overlap with GFP | For amplification of cls_TM fragment |
| GFP/(cls_TM)/5’ | tgcgcacactAGTAAAGGAGAAGCACTTTTC | lower case for overlap with cls_TM | For amplification of GFP |
| GFP/(pBOMB)/3’ | tttgaatggtcgaccggtacTTATTTGTATAGTTCATCCATGCCATG | lower case for plasmid overlap construction | For amplification of GFP |
| ct284 cls qPCR F | CACCTGTTGGCCGCTATTA | | Transcript analysis |
| ct284 cls qPCR R | GCGCTGAAGAACTGTGGATA | | Transcript analysis |

| gBlock name | Sequence | Features | Usage |
|---|---|---|---|
| c11orf83_6xH (pBOMB) | aaagaggagaaaggatctgcATGGACTCACTCCGTAAGATGTTAATATCCGTCGCTATGCTTGGAGCTGGTGCAGGTGTTGGCTACGCATTACTAGTTATCGTGACCCCCGGCGAACGCAGAAAACAGGAAATGCTCAAGGAAATGCCTTTACAGGACCCACGTTCAAGAGAAGAAGCGGCCCGAACGCAACAGTTACTCTTAGCAACCTTACAGGAAGCTGCTACAACACAGGAGAATGTTGCCTGGAGAAAAAATTGGATGGTAGGTGGGGAAGGCGGTGCAGGTGGAAGATCCCCG**CACCATCACCATCACCAT**TAAgtaccggtcgaccattcaaa | lower case for plasmid overlap, bolded is 6xH | Insert codon-optimized c11orf83_6xH into pBOMB |
| cls_TM_c11orf83_6xH (pBOMB) | aaagaggagaaaggatctgcATGAAAATGGCTTTTTTACGGAAAATATTTGTATTTGTAGCTTGTGTTGTCTCGTTGAATGGTTTTGCGCACACTGACTCACTCCGTAAGATGTTAATATCCGTCGCTATGCTTGGAGCTGGTGCAGGTGTTGGCTACGCATTACTAGTTATCGTGACCCCCGGCGAACGCAGAAAACAGGAAATGCTCAAGGAAATGCCTTTACAGGACCCACGTTCAAGAGAAGAAGCGGCCCGAACGCAACAG | lower case for plasmid overlap, underlined sequence is cls_TM, bolded is 6xH | Insert codon-optimized cls_TM_c11orf83_6xH into pBOMB |

| | | | |
|---|---|---|---|
| | TTACTCTTAGCAACCTTACAGGAAGCTGCTACAACACAGGAGAATGTTGCCTGGAGAAAAAATTGGATGGTAGGTGGGGAAGGCGGTGCAGGTGGAAGATCCCCG**CACCATCACCATCACCAT**TAAgtaccggtcgaccattcaaa | | |
| TM_c11orf83_6xH (pBOMB) | gatctaaagaggagaaaggatctgcATGTTAAAATTTATTCTACGTCGCTGTCTGGAAGCGATTCCGACGCTATTTATTCTTATTACTATTTCGTTCTTTATGATGCGCCTCGCGCCGGGAAGCCCTTTTACC*GGTGGATCA*TTACTAGTTATCGTGACCCCCGGCGAACGCAGAAAACAGGAAATGCTCAAGGAAATGCCTTTACAGGACCCACGTTCAAGAGAAGAAGCGGCCCGAACGCAACAGTTACTCTTAGCAACCTTACAGGAAGCTGCTACAACACAGGAGAATGTTGCCTGGAGAAAAAATTGGATGGTAGGTGGGGAAGGCGGTGCAGGTGGAAGATCCCCG**CACCATCACCATCACCAT**TAAgtaccggtcgaccattcaaatatgt | lower case for plasmid overlap, underlined sequence is oppB TM from *E. coli*, italicized sequence is GGS linker, bolded is 6xH, c11orf83 sequence lacks any predicted membrane-targeting motifs (ΔN23) | Insert codon-optimized TM_c11orf83_6xH into pBOMB |
| Ec_oppB_TM (pBOMB) | gatctaaagaggagaaaggatctgcATGTTAAAATTTATTCTACGTCGCTGTCTGGAAGCGATTCCGACGCTATTTATTCTTATTACTATTTCGTTCTTTATGATGCGCCTCGCGCCGGGAAGCCCTTTTACC*GGTGGATCA*ggcatggtctctaagggcgaggaaga | lower case for plasmid overlap, underlined sequence is oppB TM from *E. coli*, italicized sequence is GGS linker | Insert *E. coli* OppB 1st TM domain into pBOMB |


1 Bauler, L. D. & Hackstadt, T. Expression and targeting of secreted proteins from Chlamydia trachomatis. *J Bacteriol* **196**, 1325-1334, doi:10.1128/JB.01290-13 (2014).

2. Lee, J., Cox, J.V., & Ouellette, S. P. Critical role for the extended N-terminus of chlamydial MreB in directing its membrane association and potential interaction with divisome proteins. *J Bacteriol* **202**, e00034-20 (2020).

3. Wood, N. A., Blocker, A. M., Seleem, M. A., Conda-Sheridan, M., Fisher, D. J., & Ouellette, S. P. The ClpX and ClpP2 orthologs of *Chlamydia trachomatis* perform discrete and essential functions in organism growth and development. *mBio* **11**, e02016-20 (2020).

4 Ouellette, S. P., Gauliard, E., Antosova, Z, & Ladant, D. A Gateway®-compatible bacterial adenylate cyclase-based two-hybrid system. *Environ Microbiol Rep* **6**: 259-267 (2014).
